# Neuronal activity promotes axonal node-like clustering prior to myelination and remyelination in the central nervous system

**DOI:** 10.1101/2024.03.16.585168

**Authors:** Rémi Ronzano, Clément Perrot, Elisa Mazuir, Melina Thetiot, Marie-Stéphane Aigrot, Paul Stheneur, François-Xavier Lejeune, Bruno Stankoff, Catherine Lubetzki, Nathalie Sol-Foulon, Anne Desmazières

## Abstract

Nodes of Ranvier ensure the fast saltatory conduction along myelinated axons, through their enrichment in voltage-gated sodium and potassium channels. We and others have shown that node-like cluster assembly can occur before myelination. In multiple sclerosis, demyelination is associated with node of Ranvier disassembly, but node-like reassembly can occur prior to remyelination. Given the crucial role of neuronal activity in inducing (re)myelination, we asked whether neuronal activity could regulate node-like clustering.

We show that node-like clustering is promoted by neuronal activity and decreased when excitatory glutamatergic receptors are inhibited. Altering glutamatergic neurotransmission leads to the downregulation of Nav1.1 expression, which we show to be critical for node-like clustering. Neuronal activity also promotes node-like clustering in remyelination. As node-like clusters modulate conduction velocity and myelination initiation along axons, we propose that activity-dependent node-like clustering could modulate neuronal network establishment, as well as myelination regulation and patterning during development, plasticity and repair.

## INTRODUCTION

Myelination ensures the fast saltatory conduction along axons through the alternance of axonal segments electrically insulated by myelin and the nodes of Ranvier, which are the small amyelinic excitable domains allowing the propagation of action potentials. In diseases such as multiple sclerosis, myelin loss, associated with nodal-protein diffusion along axons, can lead to functional deficits and neuronal damage. The reorganization of myelinated fibers through remyelination can restore altered functions, as well as ensure neuroprotection ^1,2^. Nodal structures were classically described to assemble concomitantly with myelination ^3–6^. An alternative mechanism of assembly of node-like clusters (or prenodes) has been characterized prior to myelin deposition in the central nervous system by our team and others and shown to depend on both oligodendroglial secreted cues and neuronal intrinsic factors^7–13^. Node-like clusters have been described along the axon of various neuronal subtypes, including parvalbumin and somatostatin GABAergic neurons of the hippocampus, cerebellar Purkinje cells and retinal ganglion cells ^2,7–13^. These structures are observed in all the vertebrate species examined, namely zebrafish, mouse, rat, and human ^2,13,14^. Interestingly, they can form along both glutamatergic or GABAergic neurons, and appear to be more specifically associated with long-projecting or highly complex axons ^2^. In multiple sclerosis brain, similar structures have been reported in remyelinating lesions ^14^.

Node-like clusters contain the main markers of nodes of Ranvier, such as Nav channels (Nav1.1, Nav1.2, with Nav1.6 expression being more sparse or absent), associated to β1Nav and β2Nav subunits, Nfasc186, NrCAM and Ankyrin G (AnkG) ^7–10,12^. A strong diffuse expression of Caspr and Kv1.2 has been observed along axons with node-like clusters (^7^ and our unpublished results), but whether – and which – nodal potassium channels could be clustered at the node-like structures has not yet been investigated. It was however shown by patch scPatch-RNA sequencing that *KCNQ2/3* (Kv7.2/3), *KCNC1* (Kv3.1b) and *KCNK4* (TRAAK1) are expressed in node-like forming hippocampal GABAergic neurons in culture, and it is described that all three can be found at mature nodes ^2,15^.

Functionally, node-like clusters have proved to accelerate axonal conduction velocity. They further participate in heminodal and mature node assembly and guide myelination initiation at their direct vicinity ^8,9,13,16^. In the zebrafish, a subset of node-like clusters is thought to predefine mature nodes of Ranvier localization ^13^, suggesting that they could be landmarks for axonal domain and myelination pattern organization. It is thus of importance to better understand the mechanisms regulating node-like cluster formation in development and disease.

Neuronal activity is known to play a key role in the induction of myelination ^17–25^. It has been shown to modulate remyelination under pathological conditions ^26–28^ and to participate in adaptative myelination, which allows the fine tuning of neuronal networks in different areas of the central nervous system (CNS) to promote coordinated activity and memory consolidation ^29–32^. Of note, these previous studies explored how neuronal activity influences myelination and remyelination per se, but did not unravel its effects on axonal microdomain organization.

Here, we have investigated in various neuronal subpopulations whether neuronal activity regulates the assembly of node-like clusters prior to myelin deposition. We first have modulated neuronal activity and glutamatergic excitatory neurotransmission in mixed hippocampal cultures, where node-like clusters form along the axons of GABAergic neurons. In this context, neuronal activity promotes node-like cluster assembly, whereas the inhibition of glutamatergic receptors leads to a reduction of node-like clustering. Glutamatergic signaling inhibition is further associated to a decrease of the expression of some nodal proteins, including the voltage-gated sodium channel isoform Nav1.1, which we show to be critical for node-like clustering. The role of neuronal activity in node-like cluster assembly has further been confirmed using chemogenetics and optogenetics stimulations *ex vivo* in organotypic cultures of mouse cerebellar slices. In addition, neuronal activity also promotes node-like clustering in remyelination ex vivo, as well as in vivo, further strengthening physiological relevance.

## RESULTS

### Neuronal activity modulates node-like clustering along the axon of GABAergic neurons in vitro

In mixed hippocampal cell cultures, node-like clustering is first observed around 14 days in vitro (DIV) on a subset of GABAergic neurons, with the percentage of GABAergic neurons with node-like clusters increasing over time ^8^. In these cultures, neurons are electrically active, with GABAergic neurons firing action potentials (APs) of short duration and receiving high frequencies of glutamatergic excitatory synaptic events ^15^.

To assess whether node-like clustering could be regulated by neuronal activity, we first used AAV transduction of mixed hippocampal cell cultures to express the DREADDs receptors hM3D(Gq)-mCherry (activation) or hM4D(Gi)-mCherry (inhibition) under the control of the pan-neuronal human Synapsin promoter. We then assessed the percentage of GAD+/mCherry+ neurons with node-like clusters, following treatment with the DREADDs activator N-Clozapine (CNO, 0.5μM) from 8 DIV until fixation of the cultures, compared to control condition (Ctrl, Figure 1A).

**Figure 1.**
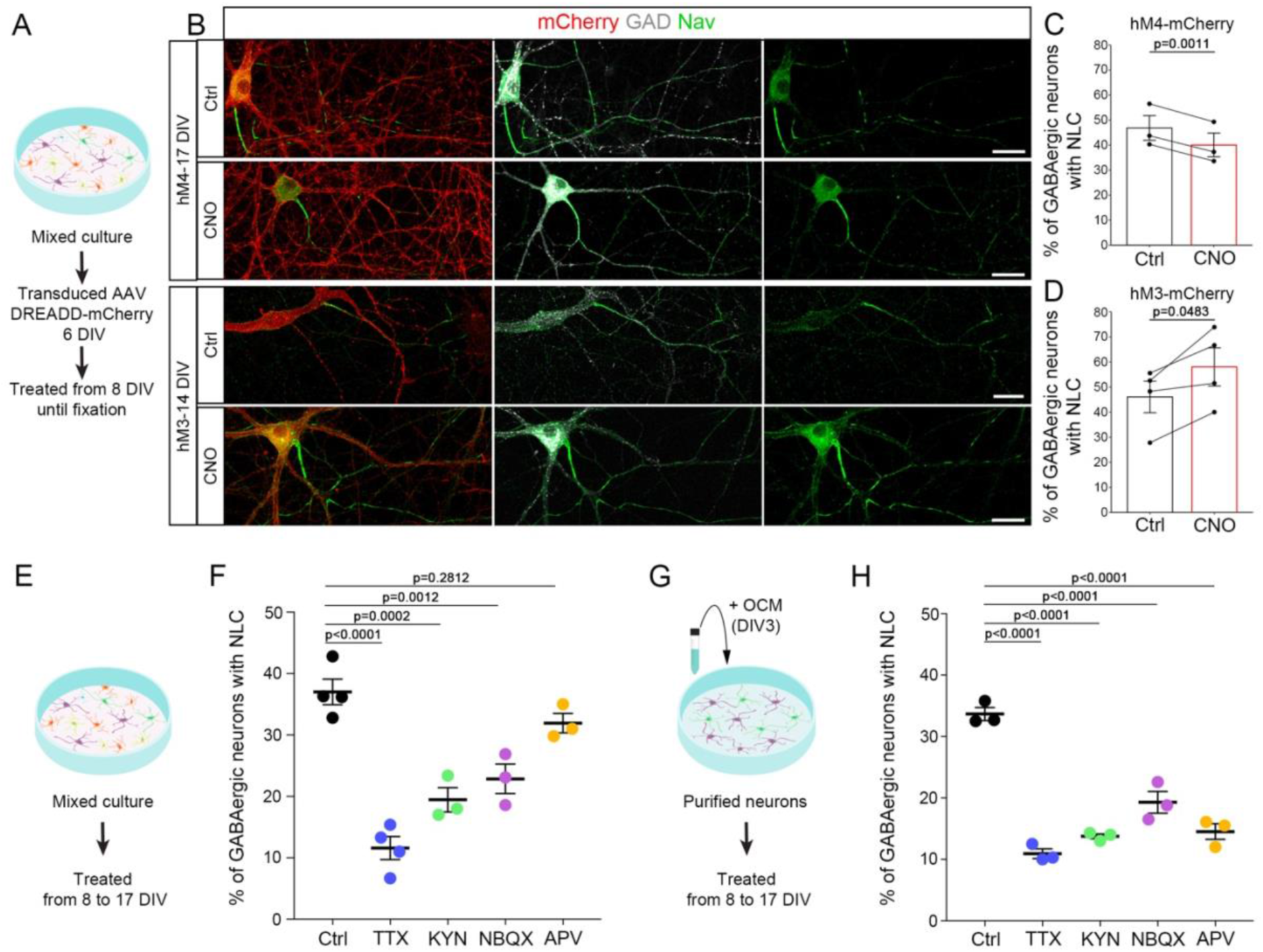
Node-like clustering along the axon of GABAergic neurons is modulated by neuronal activity and glutamatergic inputs. (A) Neuronal activity was modulated by expression of the DREADDs receptor hM4D(Gi)-mCherry or hM3D(Gq)-mCherry in neurons in mixed hippocampal cultures, followed by the addition of the DREADDs ligand N-Clozapine (CNO, 0.5μM) to the cultures, while control cultures were treated with DMSO only. (B) Illustration of GABAergic neurons (GAD67+, white) expressing DREADDs receptor hM4D(Gi)-mCherry or hM3D(Gq)-mCherry (mCherry+, red) with node-like clusters along their axons (Nav, green) in control condition or following CNO treatment. (C, D) Percentage of transduced hM4D(Gi) (C) and hM3D(Gq). (D) GAD+ neurons with node-like clusters in control condition or following CNO treatment. The values plotted individually correspond to the percentage obtained for the different experiments (hM4D(Gi), n=3 experiments; hM3D(Gq), n=4 experiments). (E, G) Mixed hippocampal cultures (E) or purified neuron cultures supplemented with oligodendrocyte conditioned medium (OCM, G) were treated with TTX (0.1µM) or glutamatergic antagonists (kynurenic acid, KYN, 1mM; NBQX, 10µM or APV, 100µM). (F, H) Percentage of GAD+ neurons with node-like clusters following treatment by TTX, KYN, NBQX or APV in mixed primary culture (F) and purified neuron culture treated with OCM (H). (C, D) Two-sided paired t-tests. (F, H) One-way ANOVA followed by Dunnett’s multiple comparison test. Scale bars: 30 μm.

The inhibition of neuronal activity mediated by hM4D(Gi)-mCherry led to a small but significant decrease in GAD+ mCherry+ neurons with node-like clusters (Ctrl=46.9 ± 4.9%; CNO=40.1 ± 4.7%; Figure 1B, C), whereas the activation of neuronal activity mediated by hM3D(Gq)-mCherry led to a 25% increase in the population of GAD+ mCherry+ neurons with node-like clusters (ctrl=46.1 ± 6.3%; CNO=58.0 ± 7.6%; Figure 1B, D). These results show that neuronal activity promotes node-like clustering in GABAergic neurons in vitro.

### The pharmacological inhibition of activity and glutamatergic signaling leads to reduced node-like clustering in vitro

The DREADDs strategy is based on GPCR (G-Protein coupled receptors), which upon activation can also lead to the activation of Gi and Gq canonical pathways, in addition to neuronal activity modulation ^33,34^. To confirm the direct role of neuronal activity on the regulation of node-like clustering, we used tetrodotoxin (TTX), a toxin blocking Na+ ion flux through the Nav channel, thereby preventing AP generation and propagation. The addition of TTX (0,1µM) in mixed hippocampal cultures from 8 DIV to 17 DIV (Figure 1E) led to a 69% reduction in the percentage of GABAergic neurons with clusters compared to control condition (Ctrl: 37.0 ± 2.1 %; TTX: 11.6 ± 1.9 %; Figure 1 F). To assess whether the modulation of node-like clustering could be due to non-autonomous cellular mechanisms mediated through the glial cells, and in particular to the oligodendrocytes present in the culture, we repeated these experiments on purified hippocampal neurons treated with oligodendrocyte conditioned medium (OCM; ^8^). A similar effect of TTX was observed in this condition compared to mixed hippocampal cultures, with a strong decrease in node-like clustering (Ctrl: 33.7 ± 1.1 %; TTX: 10.9 ± 0.8 %; Figure 1G, H). Thus, neuronal activity can modulate node-like clustering by directly regulating neuronal properties.

We next determined the effect of excitatory glutamatergic signals on node-like cluster formation. We used different antagonists of glutamatergic receptors, i.e. kynurenic acid (KYN), which inhibits AMPA, Kainate and NMDA receptors, as well as NBQX and APV, which are selective inhibitors of AMPA/Kainate and NMDA receptors respectively (Figure 1E). Adding KYN induced a 2-fold reduction in clustering activity in mixed cultures and OCM treated cultures (Mixed cultures: Ctrl: 37.0 ± 2.1 %; KYN: 19.4 ± 2.0 %; OCM: Ctrl: 33.7 ± 1.1 %; KYN: 13.8 ± 0.4 %; Figure 1F, H). Similar results were observed using NBQX (Mixed cultures: 22.9 ± 2.4 %; OCM: 19.3 ± 1.8 %), whereas APV only induced a significant reduction of clustering activity on purified hippocampal neurons treated with OCM (14.5 ± 1.3 %). These data show that neuronal glutamatergic receptors participate in node-like clustering modulation.

### The inhibition of glutamatergic neurotransmission alters the expression of specific nodal proteins in hippocampal GABAergic neurons

We first asked whether glutamatergic signals could impact node-like clustering through the modulation of nodal protein expression. For this purpose, we quantified the percentage of GABAergic neurons expressing various proteins known to be present at the axon initial segment (AIS) and at node-like clusters in GABAergic neurons, comparing mixed hippocampal cultures treated with kynurenic acid (KYN) with control condition.

We first observed that AnkG and PanNav stainings are always present in GABAergic neurons, at least at the AIS, independently of the condition considered (Figure 2A-C, G-I). GABAergic cells can express Nav1.1, Nav1.2 and Nav1.6 isoforms at node-like clusters and axon initial segments, while Nav1.1 is absent of glutamatergic hippocampal neurons^8^. We assessed the expression of Nav1.1, as well as of Neurofascin (Nfasc), Nav subunits β1Nav and β2Nav and the potassium channels KCNQ2/Kv7.2 and Kv3.1b in GABAergic neurons in control condition and following KYN treatment.

**Figure 2.**
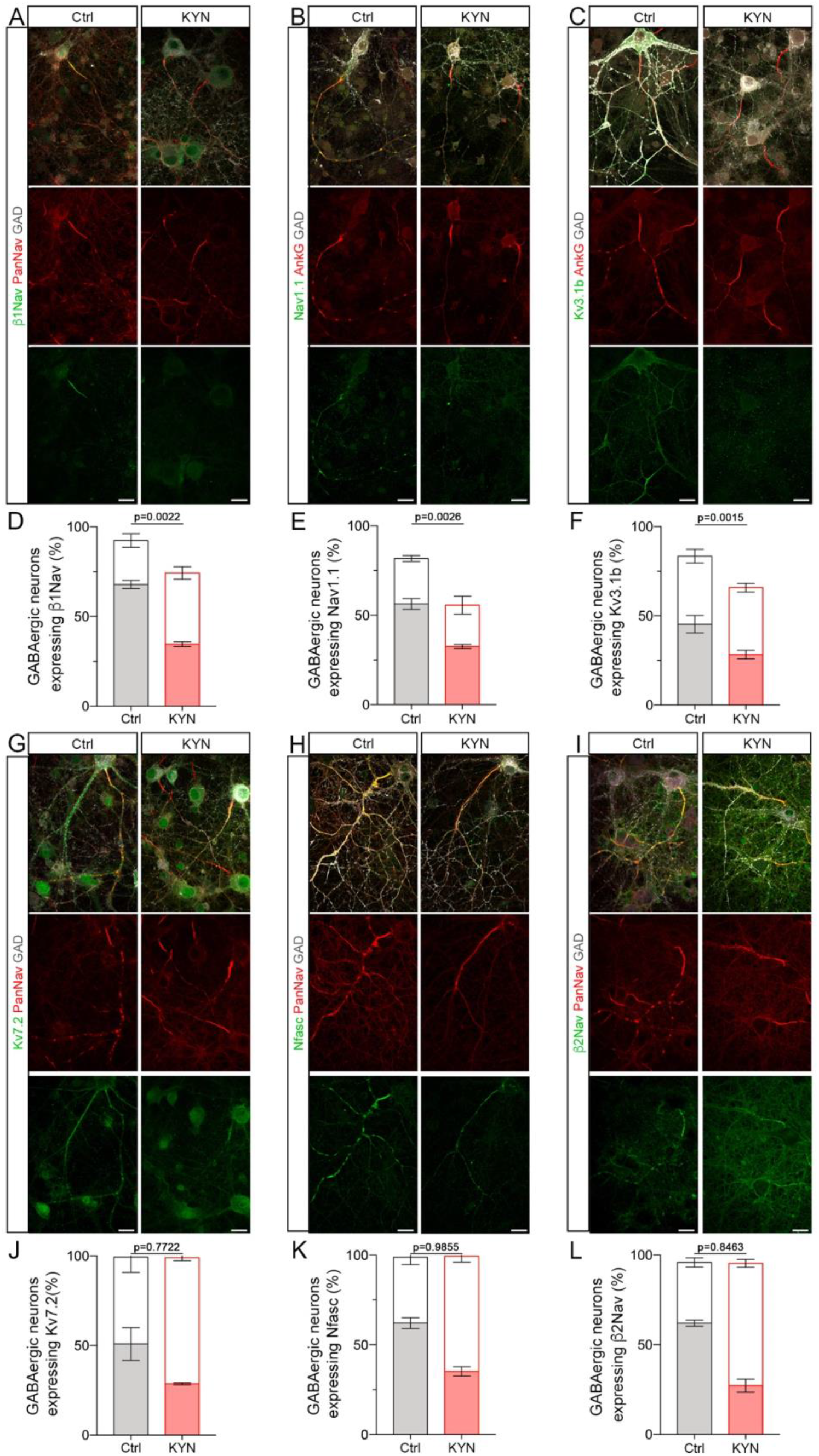
Glutamatergic inputs modulate specific nodal marker expression in GABAergic neurons. The inhibition of glutamatergic inputs in mixed hippocampal cultures by the addition of a glutamatergic antagonist (kynurenic acid, KYN, 1mM) affects the expression in GABAergic neurons (GAD+, white) of β1Nav (A, D), Nav1.1 (B, E) and Kv3.1b (C, F), in green, Kv7.2 (G, J), Nfasc (H, K) and β2Nav (I, L), all three in green, are still expressed in the vast majority of GABAergic neurons. Of note, at the timepoint considered, in control condition, Kv7.2 was restricted to the axon initial segment and node-like clusters when present. Kv3.1b, though strongly expressed in some GAD+ neurons, was not restricted to, nor enriched at axonal domains. (D-F, J-L) Percentages of GAD+ neurons expressing the nodal marker of interest (percentage of cells with node-like clusters in dark and of cells without clusters in white) in control condition or following treatment by KYN. Unpaired t-test. Scale bars: 20 μm.

While β2Nav, Nfasc and KCNQ2 were expressed in the vast majority of the GABAergic neurons in control and KYN condition, the percentages of GABAergic neurons expressing Nav1.1, β1Nav and Kv3.1b were significantly decreased following KYN treatment (Figure 2A-F; Nav1.1: Ctrl = 81.7 ± 2.9 %; KYN = 55.7 ± 4.5 %; β1Nav: Ctrl = 92.4 ± 1.7 %; KYN = 67.7 ± 3.1 % and Kv3.1b: Ctrl = 83.4 ± 2.0 %; KYN = 65.8 ± 2.5 %; Figure 2F).

The decreased expression of Nav1.1, β1Nav and Kv3.1b was linked to a loss of expression of these proteins in a subpopulation of GABAergic neurons deprived of node-like clusters, while their expression was preserved in the vast majority of GABAergic neurons with clusters (Figure S1A-C).

We further assessed whether the axonal transport of nodal markers could be affected following glutamatergic signal alteration. We expressed the nodal markers β1Nav or β2Nav coupled to mCherry by transfection of mixed hippocampal cultures and imaged the axonal transport of the mCherry+ puncta by videomicroscopy at 16-17 DIV as previously done ^9^. We then generated kymographs representing puncta axonal trajectories along time (Figure S1D). For both markers, the mean number of puncta per 100μm and the distribution of puncta amongst various categories (anterograde, retrograde or bidirectional movement, as well as non-moving puncta) showed no significant differences between KYN and control conditions (Figure S1E-G: number of puncta per 100 μm: β1Nav Ctrl = 89 ± 17; KYN = 93 ± 22; β2Nav Ctrl = 80 ± 19; KYN = 79 ± 21 and S1F-H: puncta distribution, see statistical table for detailed Mean+/-SEM).

These results suggest that glutamatergic inputs can modulate specifically some nodal protein expression in hippocampal GABAergic neurons, which in turn could lead to a variation of node-like cluster assembly in these cells.

### Nav1.1 is required for efficient node-like cluster assembly in GABAergic neurons in vitro

Amongst the nodal proteins observed to be modulated by glutamatergic inputs, Nav1.1 was the only one consistently expressed in all the node-like clusters observed along GABAergic axons (Figure 2B,^8,9^).

This led us to assess whether Nav1.1 expression was required to efficiently assemble node-like clusters in vitro. We thus designed two miRNA (miR 254 and miR 947) specifically targeting *SCN1A* (which encodes Nav1.1), to be compared with a control miRNA. All miRNA were coupled to emGFP expression in order to ensure the detection of the transfected neurons. We first confirmed that Nav1.1 expression was lost following the expression of the *SCN1A*-targeting miRNA in GABAergic neurons, while Nav1.1 was still expressed in GABAergic neurons expressing the control construct (Figure 3A). We then quantified the percentage of emGFP+ GABAergic neurons with node-like clusters at 17 DIV, comparing the two *SCN1A*-targeting miRNA with the control construct (Figure 3B-E), using AnkG or PanNav staining to detect the clusters. With PanNav staining, we observed a strong and significant decrease in the percentage of GABAergic cells with node-like clusters when miRNA 254 or miRNA 947 were expressed compared to the control condition (Figure B-C, 45% and 61% decrease respectively). This was confirmed by further using AnkG to quantify node-like clustering in the presence of miRNA 254 or 947 (Figure D-E, 61% and 66% decrease respectively) compared to the control miRNA.

**Figure 3.**
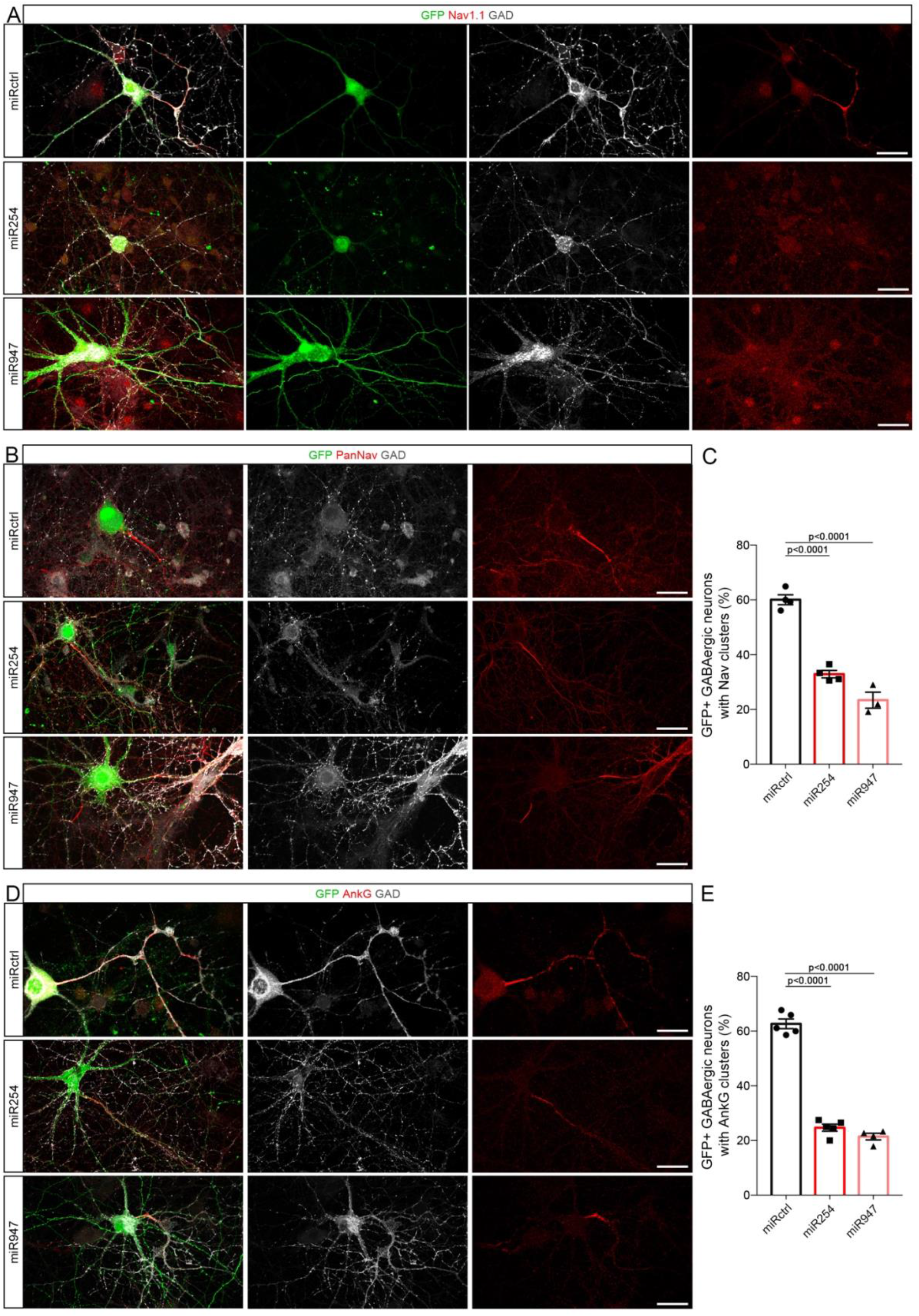
Nav1.1 downregulation leads to decreased node-like clustering. (A) Both *SCN1A*-targeting miRNA (miR254 and miR947) lead to an efficient downregulation of Nav1.1 expression (in red), compared to control miRNA (miRCtrl), in transfected GABAergic neurons (GAD+ neurons, in white, expressing emGFP, in green). (B, D) Node-like clustering is reduced in GABAergic neurons expressing *SCN1A* miRNA compared to control miRNA, as seen with Nav (B, in red) and AnkG stainings (D, in red). (C, E) Percentage of GAD+ transfected neurons with node-like clusters, assessed using PanNav (C) or AnkG (E). The percentage of neurons with node-like clusters is significantly reduced following Nav1.1 downregulation compared to control with both stainings. (C, E) One-Way ANOVA, followed by Dunnett’s multiple comparison test. Scale bars: 30 μm.

These results show that Nav1.1 is required for efficient node-like assembly and suggest that glutamatergic inputs can impact node-like clustering by modulation of Nav1.1 expression.

### Node-like clustering is modulated by neuronal activity prior to myelination in the cerebellum ex vivo

As mentioned above, node-like clusters are not restricted to hippocampal GABAergic neurons but can be transiently present prior to myelination in various subsets of neurons. We in particular observed the assembly of such structures along cerebellar Purkinje cells during developmental myelination in vivo, as well as in cultured cerebellar slices ex vivo (Figure S2A-B and D-E), with a similar structure density at the onset of myelination (Figure S2F, in vivo: 340 ± 61 per mm^2^; ex vivo: 426 ± 67 per mm^2^). To assess whether the modulation of node-like clustering by neuronal activity is a general feature, we developed two approaches to modulate neuronal activity in Purkinje cells using organotypic cultures of cerebellar slices.

We first expressed hM3D(Gq)-mCherry by AAV transduction of cerebellar slices and treated the slices with N-Clozapine at the very onset of myelination. We first confirmed that this strategy allowed a significant increase in neuronal activity by patch clamp of mCherry^+^ Purkinje cells (Figure S3A). The recordings performed showed a 4-fold increase in Purkinje cell discharge frequencies following CNO addition to the medium (Ctrl = 4.5 ± 1.0 Hz; CNO = 18.4 ± 4.0 Hz; Figure S3B-C).

We next assessed whether DREADD-mediated increase in neuronal activity could modulate node-like clustering at the onset of myelination (Figure 4A-B). We indeed observed a 1.4-fold increase in the percentage of mCherry^+^ Purkinje cells with node-like clusters following the reinforcement of their firing activity (6-hour CNO treatment, 3 DIV, Ctrl = 39.7± 2.9 %; CNO = 56.2 ± 3.3 %; Figure 4C-D).

**Figure 4.**
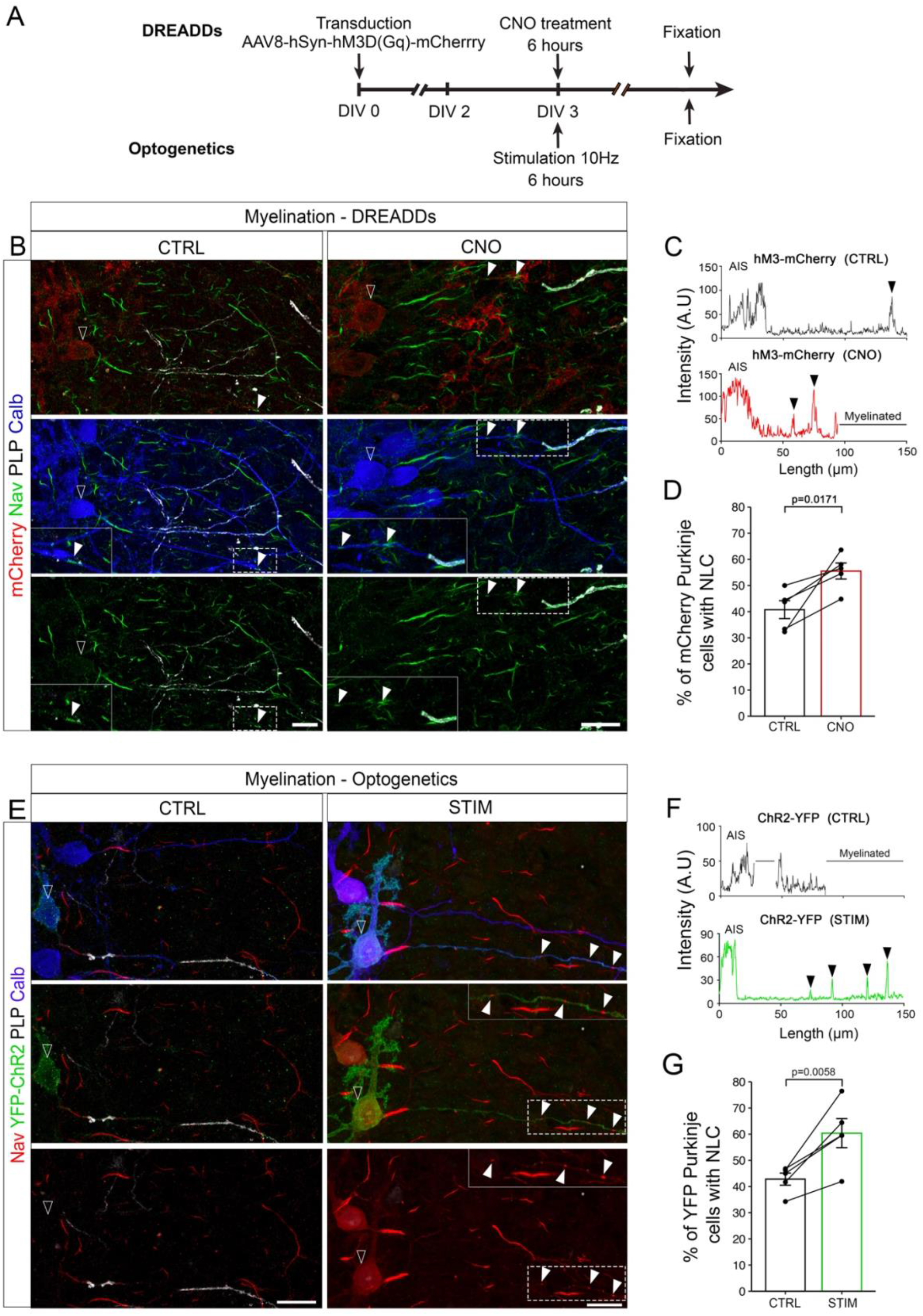
Neuronal activity enhances node-like clustering along Purkinje cells prior to myelination. For the DREADDs approach, cerebellar slices were transduced with AAV8-hSyn-hM3D(Gq)-mCherry after being generated, treated with CNO or DMSO (Ctrl) at 3 DIV for 6 hours and fixed. For the optogenetics approach, L7-ChR2-YFP mouse cerebellar slices were illuminated (470nm, STIM) or not (Ctrl) for 6 hours at 3 DIV, before being fixed. (B) Purkinje cells (Calbindin, blue) transduced with DREADDs-expressing AAV (mCherry+, red, contour arrowhead) with node-like clusters along their axon (Nav, green, filled arrowhead) in control or CNO-treated slices. (C) Plot profile of Nav staining intensity along mCherry+ Purkinje cell axons (contour arrowhead) in control (Ctrl) or CNO condition shown in B. Arrowheads show node-like clusters. (D) Percentage of mCherry+ Purkinje cells with node-like clusters following CNO (0.5 μM) or DMSO treatment (Ctrl). (E) Purkinje cells (Calbindin, Blue) expressing ChR2 (YFP, Green, white contour arrowhead) with node-like clusters along their axon (Nav, red, white filled arrowhead). Arrowheads indicate node-like clusters. (F) Plot profile of Nav staining intensity along YFP+ Purkinje cell axons in control and stimulated condition shown in E. (G) Percentage of YFP+ Purkinje cells with node-like clusters following six hours of stimulation (470nm, STIM) or in control condition (CTRL). (D) n = 5 animals, Paired t test. (G) n=5 animals, Paired t-test. (B, E) Scale Bar: 20 μm.

To confirm these results and reinforce the neuronal specificity of our approach, we further developed a custom-designed optogenetics set-up allowing the controlled illumination at a wavelength of 470nm of cerebellar slices over time (See Material and Method section for detailed description). We used this set-up to stimulate organotypic cultures of cerebellar slices generated from L7-ChR2-YFP mice ^35^, which allows to specifically modulate Purkinje cell activity though the expression of a channelrhodopsin (ChR2) under control of the L7/pcp2 promoter. We first assessed the percentage of Purkinje cells expressing ChR2-YFP in the slices at the onset of myelination (Figure S3D-F, around 80% of YFP expressing Purkinje cells in the folia analyzed in our study). We next used an illumination pattern of 10ms pulses at 10Hz, to induce a significant increase in Purkinje cells firing while staying within their physiological firing range (^36^; Ctrl: 1.3 ± 0.7 Hz; Stimulated: 15.2 ± 2.9 Hz; Figure S3G-I).

We then assessed whether an increase in neuronal activity induced by optogenetics in Purkinje cells could modulate their node-like clustering at the onset of myelination (6-hour stimulation, 3 DIV, Figure 4A and E) and confirmed this led to a significant increase in the percentage of YFP+ Purkinje cells with node-like clusters in stimulated compared to control cells (Ctrl: 43.0 ± 1.8 %; Stimulated: 60.0 ± 4.4 %; Figure 4F-G).

Finally, we assessed whether altering glutamatergic input signaling could reduce node-like clustering along Purkinje cell axon at the onset of myelination, as previously observed for hippocampal GABAergic neurons. Organotypic cerebellar slices were exposed to kynurenic acid (KYN) and node-like clustering was quantified in Purkinje cells (Figure S4). We observed a small, but significant decrease of Purkinje axons with node-like clusters following KYN treatment (Ctrl = 54.2 ± 2.7 %; KYN = 46.0 ± 2.5 %; Figure S4B), confirming the observations made in vitro.

### Node-like clustering depends on neuronal activity during remyelination in the cerebellum ex vivo

Organotypic cerebellar slice cultures myelinate spontaneously, with the vast majority of Purkinje cell axons being myelinated, which allows to induce demyelination in these cultures using lysophosphatidylcholine (LPC) and study spontaneous remyelination of the demyelinated fibers ex vivo ^37,38^. Node-like clustering was observed in the demyelinated slices just prior to remyelination, with a similar density to the one observed ex vivo during myelination (Figure S2C, F).

We took advantage of this approach to next assess whether neuronal activity could further modulate node-like clustering during remyelination. Neuronal activity was increased using the DREADDs approach at the onset of remyelination (6-hour CNO treatment at 10 DIV, Figure 5A-B). We observed a 1.3-fold increase of the percentage of mCherry^+^ Purkinje cells with node-like clusters following the reinforcement of their activity (Ctrl = 40.6 ± 2.5 %; CNO = 51.6 ± 2.6 %; Figure 5C-D).

**Figure 5.**
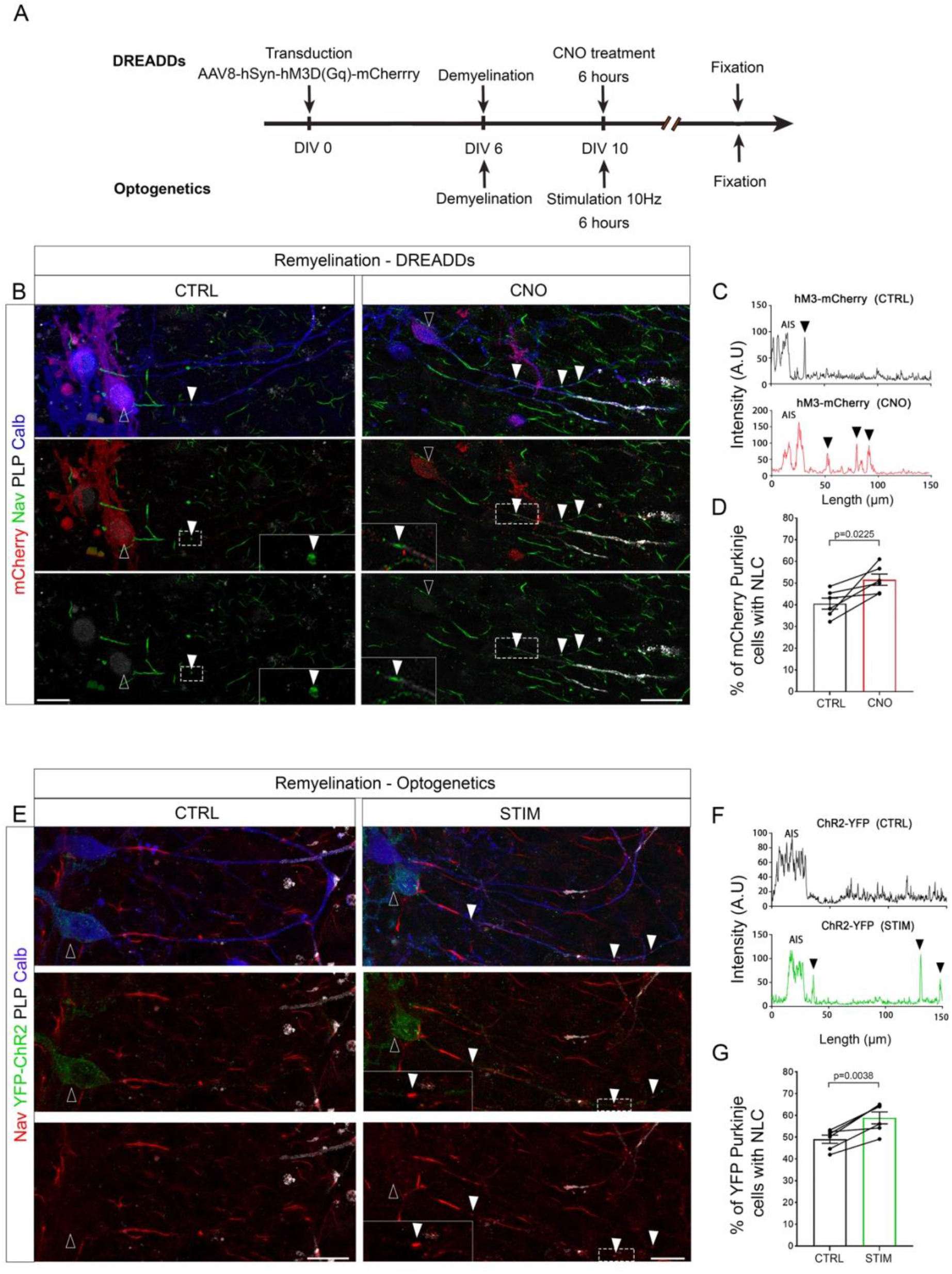
Neuronal activity enhances node-like clustering along Purkinje cells prior to remyelination. For the DREADDs approach, cerebellar slices were transduced with AAV8-hSyn-hM3D(Gq)-mCherry after being generated, demyelinated at 6 DIV (LPC, 0.5mg/mL, overnight treatment). They were next treated with CNO (0.5 μM) or DMSO (Ctrl) at DIV10 and fixed 6 hours later. For the optogenetics approach, L7-YFP-ChR2 mouse cerebellar slice were demyelinated at 6 DIV (LPC, 0.5 mg/mL) overnight treatment) and illuminated (470nm, STIM) or not (Ctrl) for 6 hours at 10 DIV, before being fixed. (B) Purkinje cells (Calbindin, blue) expressing hM3D(Gq)-mCherry (mCherry+, red, contour arrowhead) with node-like clusters along their axon (Nav, green, filled arrowheads) in control or CNO-treated slices. (C) Plot profile of Nav staining intensity along mCherry+ Purkinje cell axons in control (Ctrl) or CNO condition in B. Arrowheads indicate node-like clusters. (D) Percentage of mCherry+ Purkinje cells with node-like clusters following CNO (0.5 μM) or DMSO treatment (Ctrl). (E) Purkinje cells (Calbindin, blue) expressing ChR2 (YFP+, green, contour arrowhead) with node-like clusters (Nav, red, filled arrowheads) along their axon. (F) Plot profile of Nav staining intensity along YFP+ Purkinje cell axons from E, in the control and activated condition. Arrowheads indicate node-like clusters. (G) Percentage of YFP+ Purkinje cells with node-like clusters following six hours of stimulation (470nm, STIM) or in control condition (CTRL). (D, G) Each point corresponds to one animal. (D) n=6, Paired t test. (G) n=6, Paired t-test. (B,E) Scale Bar: 20 μm.

We next used the optogenetics stimulation previously described (6-hour stimulation at 10Hz, 10 DIV, Figure 5A and E) and confirmed this led to a significant increase in the percentage of YFP+ Purkinje cells with node-like clusters in illuminated compared to control cells (Ctrl = 49.1 ± 1.9 %; Stimulated =58.9 ± 2.7%; Figure 5F-G). Altogether, these results show that neuronal activity can promote node-like clustering at the onset of remyelination.

### Node-like cluster reassembly is modulated by neuronal activity during remyelination in vivo

We found that node-like clusters are formed along the dorsal tract of mouse spinal cord during remyelination in vivo (Figure 6). We next used an *in vivo* demyelinating mouse model, corresponding to a focal injection of LPC in the dorsal funiculus of adult mouse spinal cord (as described in ^39^), to investigate the effect of neuronal activity on node-like clustering *in vivo*. To modulate neuronal activity, we used retrograde AAV expressing hM3D(Gq)-mcherry or hM4D(Gi)-mcherry and targetted the supraspinal neurons sending descending projections to the spinal cord via the dorsal tracts ^40^. We activated the DREADD receptors 7.5 days post LPC injection (after the peak of demyelination, see methods) by CNO intraperitoneal injection, with control animals receiving NaCl 0.9% in place. We validated the expression and activation of the DREADD receptors *in vivo* by assessing cFos expression (immediate-early gene which expression is activated by neuronal activity, ^41^) in mCherry expressing neurons (motor cortex; Figure S5A-B). We observed a significant increase of cFos expression following hM3D(Gq)-mCherry activation, while the number of cFos positive cells was reduced following hm4D(Gi)-mCherry activation (Figure S5C-D respectively). We then quantified the density of node-like clusters within the lesion at 7.5 days post LPC injection, following CNO treatment and in control condition.

**Figure 6.**
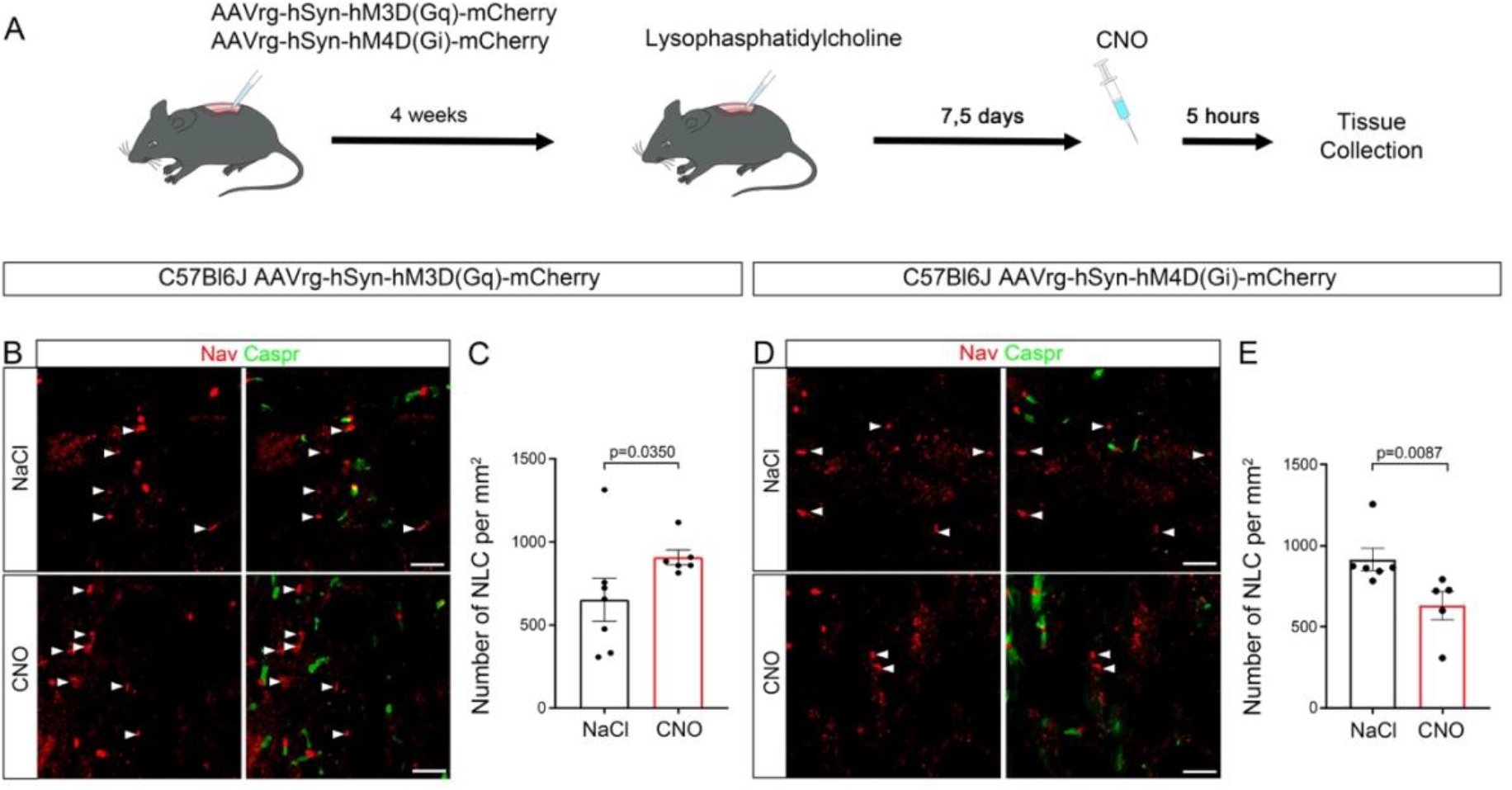
Neuronal activity increases node-like clustering in remyelinating spinal cord. (A) For the DREADDs approach, mice were transduced with a retrograde virus AAV-hSyn-hM3D(Gq)-mCherry (1μL, 2.5×10^13^ GC/ml) or AAV-hSyn-hM4D(Gi)-mCherry (1μL, 2.4×10^13^ GC/ml) in the dorsal funiculus of the spinal cord. A focal demyelination was then induced four weeks after the viral transduction, by injection of lysophosphatidylcholine (LPC, 1μL, 10mg/mL) in the same tractus to induce demyelination. Following the peak of demyelination, N-Clozapine Oxide (CNO) or NaCl (Control) was injected intraperitoneally (0.1mg) at 7.5 DPI, before perfusion of the animals. (B) Representative images of nodal clusters within the lesion of mice expressing hM3D(Gq)-mCherry. (C) Quantification of the number of node-like clusters in the lesion following CNO or NaCl injection in mice expressing of hM3D(Gq)-mCherry. (D) Representative images of nodal structures within the spinal cord lesion of mice expressing hM4D(Gi)-mCherry. (E) Quantification of the number of node-like clusters in the lesion of mice expressing hM4D(Gi)-mCherry. (B, D) Filled white arrowheads show node-like clusters (Nav, red) without paranodal staining surrounding them (Caspr, green) in CNO and NaCl condition. (C, E) Each point corresponds to one animal, (C) n=7 for NaCl and n=6 for CNO, Mann-Whitney test. (E) n=6 for NaCl and n=5 for CNO, Mann-Whitney test. (B, E) Scale Bar: 10 μm.

Following neuronal activity induction, in the case of hM3D(Gq) receptor, we observed a 1.4-fold increase of node-like cluster density in CNO treated vs control animals (Ctrl = 652 ± 129 clusters per mm^2^; CNO = 907 ± 44 clusters per mm^2^, Figure 6B-C). In a second batch of experiments, we inhibited neuronal activity at a similar timepoint post LPC injection by activation of hM4D(Gi) receptor, which led to a decrease of node-like cluster density in CNO treated animals compared to controls (Ctrl = 914 ± 69 clusters per mm^2^; CNO = 630 ± 44 clusters per mm^2^, Figure 6D-E).

These results show that neuronal activity also promotes node-like cluster assembly in remyelination *in vivo*.

## DISCUSSION

In this work, we show using *in vitro* to *in vivo* models and a range of approaches going from pharmacological treatments to optogenetics that neuronal activity modulates node-like clustering prior to myelination in various neuronal subtypes and present evidence of a similar regulatory process during remyelination, suggesting this could be a general feature of node-like cluster assembly.

### Neuronal activity and glutamatergic excitatory signals can modulate node-like clustering through a neuronal intrinsic mechanism

Neuronal activity and glutamatergic inputs are known to modulate the oligodendroglial lineage ^42–45^, which promotes node-like clustering through secreted factors (for review, ^2^). Astrocytes can further express NMDA receptors ^46^. To address the possibility of an indirect effect of neuronal activity on node-like clustering through glial cell modulation, we used purified hippocampal neuron cultures supplemented with oligodendrocyte conditioned medium (OCM) in parallel of mixed hippocampal cultures to study the effect of glutamatergic signaling inhibition. Using this approach, we still observed a modulation of node-like clustering while inhibiting glutamatergic inputs, confirming a cell-autonomous modulation of node-like clustering in GABAergic neurons. Thus, neuronal activity can regulate node-like clustering through a direct modulation of intrinsic neuronal properties, though an additional role of glial cells in response to neuronal activity glutamatergic inputs cannot be excluded. Kaplan and colleagues had previously shown that modulating activity and glutamatergic inputs pharmacologically in RGC cultured with OCM had no effects on cluster spacing ^11^. The results regarding clustering per se were somehow contradictory, though a tendency to increased clustering was observed following glutamatergic agonists treatment ^11^.

In addition, reinforcing neuronal activity was not sufficient to promote node-like clustering in hippocampal glutamatergic neurons, which do not assemble these clusters under normal condition. This further suggests that neuronal activity modulates intrinsic properties specific to the neuronal subpopulations assembling node-like clusters. The possibility that neuronal activity can modulate the characteristics of node-like clusters themselves (such as nodal length or density of channels) and whether this could further impact conduction velocity, as observed for mature nodes ^47–49^ remains to be addressed.

### Glutamatergic signaling modulates the expression of node-like cluster proteins

We further investigated which neuronal mechanisms could drive node-like clustering modulation by neuronal activity and glutamatergic signaling, and focused on nodal component expression modulation and axonal transport of these proteins, two mechanisms we previously showed to be involved in node-like clustering *in vitro* ^9,11^. Here, we demonstrated that the inhibition of glutamatergic transmission leads to a decreased expression of various nodal proteins, without major variation of nodal protein axonal transport. Neuronal activity is known to modulate neuronal transcriptome ^50–53^, and glutamatergic excitatory inputs were shown to participate in GABAergic interneurons post-natal maturation, at least in part through transcriptional regulation (for review ^54^). Moreover, in hippocampal cultures, GABAergic neurons with node-like clusters received higher frequencies of excitatory synaptic events and have been shown to differ in their transcriptome from GABAergic neurons without clusters and pyramidal cells ^15^.

These results are consistent with our findings. We observe that the inhibition of glutamatergic transmission leads in particular to a decreased expression of the voltage-gated sodium channel isoform Nav1.1, encoded by *Scn1a* gene. *Scna1* expression is developmentally regulated in cortical GABAergic neurons, and its expression also correlates with the maturation of neuronal electrophysiological properties in hippocampal GABAergic neurons ^15,55^. We previously reported that hippocampal GABAergic neurons with node-like clusters significantly upregulate *Scna1* expression compared to GABAergic neurons without clusters ^15^. This upregulation might participate in the very high density of Nav channels observed in fast-spiking GABAergic neurons, which may ensure fast signaling and compensate for the unfavorable morphological properties of their axons ^56,57^. Nav1.1 expression is a common feature of all the neuronal subtypes described to form node-like clusters ^8,10,11,58,59^ and in the hippocampus, its axonal expression is restricted to GABAergic cells, the only node-like forming neurons. In the present study, we completed these results by showing that inhibiting Nav1.1 expression leads to a strong reduction of node-like clustering in GABAergic neurons in vitro, although other Nav channels are still present as shown by the persistence of PanNav staining. Altogether this suggests that Nav1.1 regulation may be part of the mechanism through which neuronal activity regulates node-like clustering.

Though neuronal activity is described to modulate axonal trafficking ^60,61^, we did not observe alteration of nodal marker axonal transport following the inhibition of excitatory glutamatergic neurotransmission. Neuronal activity could alternatively promote initial nodal protein targeting at specific axonal “hotspots” during node-like clustering, similarly to its regulation of glutamatergic synaptic vesicular release at axonal “hotspots” promoting myelin sheath elongation ^24^.

### An indirect modulation of myelination and neuronal networks by neuronal activity through node-like clustering in development, plasticity and repair?

Neuronal activity-dependent node-like clustering could participate in various aspects of developmental processes, including neuron physiology and neuronal network establishment. Glutamatergic signaling alteration in GABAergic neurons has been associated to neurodevelopmental psychiatric disorders but whether node-like clustering is affected in these diseases has not yet been investigated. In addition neuronal node-like clustering characterizes mostly neurons that are programmed to be subsequently myelinated ^2^, and we propose that node-like clustering could be an intermediate step of activity-dependent myelination.

The formation of node like clusters along axons prior to myelination has been shown to participate to mature node formation ^9,13,16^, some node-like clusters predefining mature nodes being thus potentially landmarks for future myelination pattern. Furthermore, these clusters may regulate myelination itself by their role in conduction modulation and myelin deposition guidance ^8,9^. For this later process, local vesicular release may be at play, as it was shown to regulate myelin sheath stabilization and growth^22–24,62^, as well as promote MBP local synthesis in oligodendrocytes at close contact of release site ^63^.

This cascade of events may affect the timing of axonal transmission, which is critical for neuronal network synchronization and myelination ^48,64–66^. It would thus be of particular interest to investigate whether some non-myelinated or partially myelinated neurons can form node-like clusters in response to increased neuronal activity in adult, therefore participating in plasticity through refinement of axonal conduction and adaptive myelination ^18,67,68^.

In the present work, we show that node-like clustering is also promoted by neuronal activity prior to remyelination. In demyelinating pathologies such as multiple sclerosis, remyelination exists but is only partial and its failure may lead to neuronal loss. Node-like clustering has been previously observed in remyelinating MS plaques and in MS rodent models (Roux, Pantazou and Desmazieres, unpublished data; ^2,14,39^). The mechanism and function of node-like clustering prior to remyelination remain elusive, and whether they facilitate axonal conduction along demyelinated axons and/or promote myelin redeposition in an unfavorable environment has not yet been disentangled. In the present work, we show that node-like clustering is also promoted by neuronal activity prior to remyelination.

Extrapolating what has been suggested for developmental myelination ^13^, it can be hypothesized that node like clusters might participate in defining the pattern of remyelination along axons following demyelination ^69–71^. Of note, following cuprizone-induced demyelination, the scaffolding nodal protein βIV Spectrin can remain clustered at the site of former nodes of Ranvier ^70^, possibly serving as landmark for remyelination pattern. In contrast, the myelination pattern is not restored in some cortical areas, though the global amount of myelin is ^68^. Whether this is a defect in remyelination or an adaptation to remyelinate the most active neurons following axonal injury in cortical areas is not known. Assessing the presence and potential role of neuronal activity dependent node-like clustering in this context could give further insights in this process.

In conclusion, neuronal activity appears to support node-like clustering, which in turns modulates axonal conduction and may guide myelin deposition initiation. This could participate in the adequate establishment of neuronal networks during development, as well as myelination regulation and patterning during developmental myelination, plasticity and remyelination.

## METHODS

### Animals

The care and use of mice conformed to institutional policies and guidelines (UPMC, INSERM, French and European Community Council Directive 86/609/EEC). The following mouse strains were used in this study: C57bl6/J (Janvier Labs), L7-ChR2-eYFP (gift from Pr C. Lena, IBENS, PSL University, Paris, France, ^35^, transferred into a C57bl6/J genetic background and used after a minimum of 6 generations. Wistar pregnant rat females were purchased from Janvier Labs.

### Hippocampal mixed culture

Mixed hippocampal cultures (containing neurons and glial cells) were made from rat embryos as previously described ^8^. Briefly, pooled hippocampi of E18 rat embryos were dissociated enzymatically by trypsin treatment (0.1%; Worthington) with DNase (50 μg/ml, Worthington) for 25 min at 37°C. Following trypsin neutralization, cells were mechanically dissociated, centrifuged at 400xg for 5 min, resuspended, and seeded on polyethylenimine (Sigma) precoated glass coverslips at a density of 5.0 × 10^4^ cells per well in 24 well plates (surface of 35mm^2^ per well, TPP) or per 35 mm glass-bottom dishes (81158; Ibidi, BioValley). Cultures were maintained for 24h in a 1:1 mixture of DMEM Glutamax (31966047, ThermoFisher Scientific) with 10% fetal calf serum (10270106, ThermoFisher Scientific), penicillin-streptomycin (100 IU/mL, 15140122, ThermoFisher Scientific), and neuron culture medium (NCM). Culture medium was replaced by a 1:2 mixture of Bottenstein–Sato (BS) medium with PDGF-A (0.5%, Peprotech) and NCM, and then, half of the medium was removed every 3 days and replaced by NCM. The NCM contained neurobasal medium (21103049; ThermoFisher Scientific) supplemented with 0.5 mM L-glutamine (25030024, ThermoFisher Scientific), B27 (1×; 17504044, ThermoFisher Scientific), and penicillin-streptomycin (100 IU/mL each). The BS medium was made of DMEM Glutamax supplemented with transferrin (100 μg/mL), albumin (100 μg/mL), insulin (5 μg/mL), progesterone (60 ng/ml), putrescine (16 μg/mL), sodium selenite (40 ng/mL), T3 (40 ng/mL), and T4 (40 ng/mL), all from Sigma.

### Preparation of OCM and purified neuronal cultures supplemented with OCM

Glial cell cultures were prepared from cerebral cortices of P2 Wistar rats as described previously^72^. After meninges were removed, cortices were incubated for 35 min in papain (30 U/mL; Worthington), supplemented with L-cysteine (0.24 mg/mL, Sigma) and DNase (50 μg/mL, Worthington) in DMEM at 37°. They were then mechanically dissociated and passed through a 70-μm filter. Cells were resuspended in DMEM Glutamax with 10% FCS and 1% penicillin–streptomycin (100 IU/mL each). After 7–14 days in vitro (DIV), oligodendroglial lineage cells were purified from glial cell cultures which initially contain astrocytes and microglial cells. After cultures were shaken overnight at 230 rpm and 37 °C, overlying oligodendroglial and microglial cells could be selectively detached. Microglia were then eliminated by differential adhesion. Collected cells were incubated in dishes for 15 min. Nonadherent cells were retrieved and centrifuged in DMEM for 5 min at 400G. They were resuspended and seeded at a density of 1.5 × 10^5^/cm^2^ on polyethyleneimine-coated (PEI) dishes with BS medium and 0.5% PDGF. The cells were then incubated for two days in BS medium and then placed in NCM. The medium from these cultures was collected 48h later, filtered (0.22 μm), and stored at 4°C to be used as OCM. Purified neuron cultures (PUR) were prepared by adding anti-mitotic agents (FdU and U 5 μM) for 36h to mixed hippocampal cultures prepared as described above, starting 24 h after dissection. OCM (500 μL/well) was then added to the purified neuron cultures at 3 DIV. One-third of the medium was replaced with NCM at 7 DIV, and then twice a week.

### Plasmid constructs and cell culture transfection

The plasmids pTRIPSyn-β1NavmCherry and pTRIPSyn-β2NavmCherry used for the axonal transport study have been previously described ^9^.

The oligonucleotides used to generate the miRNA constructs targeting Nav1.1 are as follow: miR4254: Top: TGCTGTAACAGGGCATTCACAACCACGTTTTGGCCACTGACTGACGTGGTTGTATGCCCTGTTA; Bottom: CCTGTAACAGGGCATACAACCACGTCAGTCAGTGGCCAAAACGTGGTTGTGAATGCCCTGTTAC; miR4947: Top: TGCTGAATGCTTGTCACATAATCGCTGTTTTGGCCACTGACTGACAGCGATTATGACAAGCATT; Bottom: CCTGAATGCTTGTCATAATCGCTGTCAGTCAGTGGCCAAAACAGCGATTATGTGACAAGCATTC. The miRNA plasmids used for knockdown studies were generated following manufacturer’s instructions (K493600; ThermoFisher Scientific). The control miRNA construct was provided in the kit. For all the miRNA constructs used, emGFP is co-expressed with the encoded miRNA, allowing the detection of the transfected cells. Transfection of rat primary neurons was performed as previously described^8^, at 6 DIV with a total of 500ng DNA and 1.0 μl Lipofectamine 2000 reagent per well (11668019; ThermoFisher Scientific) in Opti-MEM reduced serum medium (31985062; ThermoFisher Scientific). 50ng/μl of plasmid were used per well for nodal protein expression and 400 ng/μl for miRNA, supplemented to 500ng DNA with pBlueScript vector.

### Organotypic cultures of mouse cerebellar slices

Cerebellar slice cultures were made as previously described ^37,38^. Briefly, P8-10 mouse cerebella were dissected in ice cold Gey’s balanced salt solution (G9779, Sigma) complemented with 4.5 mg/ml D-Glucose (G8769-100ML, Sigma) and 1X penicillin-streptomycin (100 IU/mL, ThermoFisher Scientific), before being cut into 250μm parasagittal slices using a McIlwain tissue chopper and placed on Millicell membrane (3 to 4 slices each per animal, 0.4 μm membranes, Merck Millipore) in 50% BME (41010026, ThermoFisher Scientific), 25% Hanks’ Balanced Salt Solution (14185-045, ThermoFisher Scientific), 25% heat-inactivated horse serum (26050088, ThermoFisher Scientific) medium, supplemented with GlutaMax (2 mM, 35050038, ThermoFisher Scientific), penicillin-streptomycin (100 IU/mL, ThermoFisher Scientific) and D-Glucose (4.5 mg/ml; Sigma). The slices from a given animal were divided on two membranes, one of the membranes being used as control, the second being treated, to minimize the potential variability and do paired experiments. Cultures were maintained at 37°C under 5% CO_2_ and medium changed every two to three days. The experiments were analyzed at 3 days in vitro (DIV, myelination) and 10 DIV (remyelination).

### Viral transduction of *in vitro* and *ex vivo* cultures

The adenoviruses used to express hM3Dq or hM4Di fused to the reporter mCherry under the control of the human Synapsin promoter, AAV8-hSyn-hM3D(Gq)-mCherry and AAV8-hSyn-hM4D(Gi)-mCherry are available commercially (gift from Bryan Roth, Addgene plasmids #50474-AAV8 and #50475-AAV8 respectively) and drove the expression of DREADDs receptors in neurons. In hippocampal mixed cultures, the transduction was performed at 6 DIV, by adding the virus at a final concentration of ∼10^9^ VP/μL, when renewing half of the medium. To activate the transduced DREAAD receptor, the cultures were treated with N-Clozapine (CNO, 0.5μM, Cayman chemical, 16882) or DMSO (700001, Cayman Chemical) for control condition from DIV 8 until fixation of the cell culture (DIV17 for hM4G(Gi)-mCherry and DIV 14 for hM3D(Gq)-mCherry. Regarding organotypic slice cultures, the transduction was performed immediately following slice preparation by addition of the AAV solution directly onto the slices placed on milicell membranes (1 μl/slice at a final concentration of 1×10^11^ and 5×10^11^ VP/μl for AAV8-hSyn-hM3D(Gq)-mCherry and AAV8-hSyn-hM4D(Gi)-mCherry respectively).

### Electrophysiology recordings

Organotypic cerebellar slices at 9 to 11 DIV were transferred to a recording chamber and continuously superfused with oxygenated (95% O2 and 5% CO2) aCSF containing (in mM): 124 NaCl, 3 KCl, 1.25 NaH2PO4, 26 NaHCO3, 1.3 MgSO4, 2.5 CaCl2, and 15 glucose (pH 7.4), all from Sigma Aldrich). Purkinje cells were visualized under differential interference contrast optics using a 63X water immersion lens (N.A. 1). Loose cell-attached voltage clamp recordings of the spontaneous firing activity of Purkinje cells were performed at 30-34°C with a borosilicate glass pipette filled with aCSF. Signals were amplified with a Multiclamp 700B amplifier (Molecular devices), sampled and filtered at 10 kHz with a Digidata 1550B (Molecular Devices). Data were acquired with the pClamp software (Molecular devices). To avoid any alteration of the spontaneous firing frequency of the cell by the patch procedure ^73^, the holding membrane potential was set to the value at which zero current was injected by the amplifier. The resistance of the seal (Rseal) was controlled and calculated every minute from the current response to a voltage step (200 ms; –10 mV). Only recordings with a Rseal in the range of 10 to 100MΩ and stable during the recording procedure were included in the analysis. To test the modulation of neuronal activity by DREADDs, recordings were performed in control and the perfusion was then switched to a bath with 0.5μM of N-clozapine (CNO) while recording the same neuron. To test optogenetics stimulation, a LED with an excitation filter 482/35 was calibrated to stimulate the field of view at 1.5mW/mm^2^ and individual neurons were successively recorded with no stimulation and with pulses of 10 millisecond at 10Hz. The mean firing rate was analyzed over 110 seconds recording time window using a threshold crossing spike detection in Clampfit (Molecular devices) and calculated as the number of action potential divided by the duration of the recording.

### Hippocampal neuron culture and organotypic culture treatments

#### Pharmacological treatment of hippocampal neuron cultures and cerebellar slices

To activate the DREADDs receptor hM3Dq or hM4Di, the culture medium was renewed with a culture medium containing 0.5μM of CNO diluted in DMSO (700001, Cayman Chemical) or a culture medium with an equivalent volume of DMSO (1μL in 1 mL of medium) in control condition. For hippocampal mixed cultures, CNO treatment started at 8 DIV and was maintained until fixation. The cerebellar slices were treated with 0.5μM CNO or DMSO (control condition) at 3 DIV (myelination) or 10 DIV (remyelination). The slices were treated 6 hours with CNO (for hM3D(Gq)) and fixed one hour after the end of the treatment to assess the effect on node-like clustering.

To study the effect of neuronal activity and glutamatergic inputs on node-like clustering in hippocampal neuron cultures, at 8 DIV half of the culture medium was renewed with an equivalent volume of NCM containing the different drugs at 2X, either tetrodotoxin (TTX, Sigma T2265-25G; final concentration 0,1µM), or glutamatergic receptor inhibitors, kynurenic acid (KYN; final concentration 1mM, Sigma), NBQX (final concentration 10µM, AbCAM) or APV (final concentration 100µM, AbCAM). Then half of the medium was renewed every three days with NCM with drug 1X until fixation at 17 DIV. To study the effect of glutamatergic inputs on node like-clustering on Purkinje cells prior myelination, kynurenic acid (KYN) was added to the culture medium directly following slice generation to a final concentration of 1mM, and the slices were fixed at 3 DIV.

#### Demyelination of organotypic cerebellar slices

To induce demyelination, for each animal, the myelinated slices were incubated 16 hours at 6 DIV in 0.5 mg/ml L-α-Lysophosphatidylcholine (LPC, 830071P, Avanti, Merck) added to fresh culture medium.

### Optogenetics stimulation of organotypic cerebellar slices

Optogenetics stimulations were performed on the organotypic cerebellar slices with a custom-designed set-up that allows stimulation with light at 470nm directly on 6-well culture plates (Desmazieres, Ronzano and Marty). Each well is illuminated by a LED (SZ-05-H, Luxeonstar), each LED being individually calibrated to allow a stimulation at a power of 1.5mW/mm^2^ on the slices. The pattern of stimulation was controlled with an Arduino (A000067, Arduino Mega 2560, Arduino, ref.782-A000067, Mouser Electronic). To stimulate Purkinje cells, we used 10 ms long pulses at 10Hz for 6 hours applied at 3 DIV or 10 DIV and fixed the slices 1 hour after the end of the stimulation to study node-like cluster formation prior to myelination or remyelination respectively. To avoid cytotoxicity, prior to stimulation, the membranes were transferred in a medium free of phenol-red consisting of: 75% DMEM (11880028, Gibco, ThermoFisher Scientific), 20% 1X HBSS (14185-045, Gibco, ThermoFisher Scientific) supplemented with HCO ^-^ (0.075 g/L final; 25080060, Gibco, ThermoFisher Scientific), 5% heat-inactivated horse serum (ThermoFisher Scientific), HEPES Buffer (10 mM final; 15630056, Gibco, ThermoFisher Scientific), D-Glucose (4.5 g/L final; G8769-100ML, Sigma), GlutaMax (2 mM final; 35050038, ThermoFisher Scientific) and penicillin-streptomycin (100 IU/mL each; ThermoFisher Scientific). The slices were fixed one hour after the end of the stimulation (effect on node-like clustering).

### In vivo Intraspinal injections

#### Viral injection

Following an intraperitoneal injection of buprenorphine (0.1mg/mL, Buprecare, Med’Vet) done 15 minutes before surgery to prevent pain, C57Bl6J females (8 weeks old, 18-22g) were anesthetized with isofluorane (3% induction, 1,5% maintenance, Vetfluran). The anesthetized animals were then placed onto a spinal stereotaxic frame and an incision was made between the thoracic and lumbar vertebrae to access the dorsal funiculus of the spinal cord. 1μL of AAVrg-hSyn-hM3D(G_q_)-mCherry (Addgene, #50474-AAVrg, 2.5×10^13^ GC/ml) or AAVrg-hSyn-hM4D(G_i_)-mCherry (Addgene, #50475-AAVrg, 2.4×10^13^GC/ml) were then injected intraspinally with a glass capillary ^40^. Following surgery, the mice were stitched and placed until recovery into a warming chamber (Vet tech solution LTD, HE011).

#### Focal spinal cord demyelination

The focal demyelination was induced 4 weeks post viral injection. The initial steps of the surgery protocol are similar to the viral injection described above. An incision is made between the thoracic and lumbar vertebrae to access the dorsal funiculus of the spine and an intraspinal injection of 1 μL of lysophosphatidylcholin diluted into NaCl 9‰ (LPC, 10mg/mL; Sigma-Aldrich L4129) is made with a glass capillary. Following surgery, the mice were stitched and placed until recovery into a warming chamber (Vet tech solution LTD, HE011). The peak of demyelination is reached around 7 days post injection, and remyelination starts from 10 days post injection.

### DREADDs receptors activation in vivo

Clozapine N-Oxide (CNO, Cayman Chemical Company) was diluted into NaCl 9‰ to a final concentration of 1mg/mL and the solution was injected intraperitoneally (5mg/kg, 2 injections with a 2-hour interval). The mice were then euthanazied with Euthasol (Centravet) and transcardially perfused with 2% PFA (Electron Microscopy Services, ThermoFisher Scientific) 5 hours after the initial CNO injection, at 7.5 days post LPC injection.

### Fixation and immunohistochemistry

#### In vitro cultured hippocampal neurons

Cell cultures were fixed at DIV14 or 17 with 4% paraformaldehyde (PFA, Electron Microscopy Services, ThermoFisher Scientific) for 10 min, or with 1% PFA for 10 min at room temperature (RT) and then incubated with methanol for 10 min at −20^°^C (for αNav staining). After fixation, cells were washed in 1× PBS, before being incubated with blocking solution (1× PBS with 5% Normal Goat Serum [50-062Z; ThermoFisher Scientific], 0.1% Triton X-100) for 15 min and with primary antibodies solutions for 2 to 3 hours at RT. Coverslips were then washed in 1× PBS and incubated with secondary antibodies solutions for 1 hr at RT. Coverslips were washed in 1× PBS and mounted on glass slides with Fluoromount G (Southern Biotech, with or without DAPI).

#### Ex vivo cultured cerebellar slices

Cerebellar slices were fixed as described before ^37^, with 4% PFA (Electron Microscopy, ThermoFisher Scientific) for 5 minutes followed by 1% PFA for 25 minutes at RT and washed in PBS. Subsequently, the slices were incubated in absolute ethanol (Sigma Aldrich) at –20°C for 20 minutes and washed in PBS. The slices were blocked for 1 hour in PBS, 5% Normal Goat Serum (50-062Z; ThermoFisher Scientific), 0.3% Triton X-100 (9036-19-5, Sigma) and incubated with primary antibodies diluted in blocking solution overnight at RT. The slices were then washed in PBS, incubated for 3 hours at RT in the dark with secondary antibodies diluted in blocking solution. The slices were washed in PBS and mounted between a glass slide and a coverslip (VWR) with Fluoromount-G (0100-01, Southern Biotech).

#### Mouse central nervous tissues fixation and collection

Adult and P10 mice were perfused with 2% PFA and the brain and spinal cord collected and post-fixed in PFA 2% for 30 minutes, washed in PBS and incubated in PBS with 30% sucrose (S0389, Sigma Aldrich) for 3 days at 4°C for cryoprotection. The tissues were then included in O.C.T (Tissue-Tek, Sakura). Using a cryostat (Leica CM 1950), the brains were cut sagittaly or coronally and the spinal cord cut longitudinally in 30μm and 20µm thick sections respectively. Sections were collected on Superfrost+ glass slides (48311-703, VWR). For immunohistochemistry, the slides were first placed in absolute ethanol at –20°C for 20 minutes. They were then incubated with a blocking solution containing PBS, 5% Normal Goat Serum and 0.2% Triton X-100 for at least 30 minutes at RT. Following PBS washes, the slides were incubated with the primary antibodies diluted in blocking solution overnight at RT and the next day with the secondary antibodies diluted in blocking solution for 2 hours in the dark at RT. The slides were mounted under a converslip (VWR) using Fluoromount with or without Hoechst (Southern Biotech), and left to dry at RT before being stored at 4°C.

### Antibodies

The following primary antibodies were used: mouse IgG2a anti-AnkyrinG (clone N106/36; 1:100, Neuromab), mouse IgG2b anti-AnkyrinG (1:75; clone N106/65, NeuroMAB), rabbit anti-AnkG (1:300, custom anti-peptide polyclonal antibody, targeted against LLERSSITMTPPASPKSN sequence, Eurogentech), mouse IgG1 anti-Pan Na_v_ (1:150-1:300, Sigma), mouse IgG1 anti-Nav1.1 (1:100, clone K74/71, NeuroMAB), rabbit anti KCNQ2/Kv7.2 (1/250, 368.103, Synaptic System), mouse IgG1 anti Kv3.1b (1:100, clone N16b8, NeuroMAB), rabbit anti-β1Nav (1:100; kindly provided by Pr P.J. Brophy, University of Edinburgh, UK), rabbit anti-β2Nav (1:250; Millipore), rabbit anti-Nfasc (pan; 1:100; Abcam), rabbit anti-Caspr (1:300, Abcam), mouse anti-Calbindin (1:500; Sigma), rabbit anti-Calbindin (1:300; Swant), rat anti-PLP (1:10; hybridoma, kindly provided by Dr. K. Ikenaka, Okasaki, Japan), chicken anti-GFP (1:250, Millipore), chicken anti-mCherry (1:1000 to 1:2000, EnCor Biotechnology), mouse IgG2a anti-GAD67 (clone 1G10.2; 1:400, Millipore), mouse IgG1 cFos (1:500; Abcam). Secondary antibodies corresponded to goat or donkey anti-chicken, mouse IgG2a, IgG2b, IgG1, rabbit, rat coupled to Alexa Fluor 488, 594, 647 or 405 from Invitrogen (1:500 to 1:1000).

### Culture and tissue imaging

#### Videomicroscopy

For axonal transport live imaging, cells were grown on 35 mm glass-bottom dishes (81158; Ibidi, BioValley), transfected as described above and imaged at DIV16-17. The culture medium was replaced by a culture medium without phenol red prior to imaging. During live imaging, dishes were placed in a temperature-controlled imaging chamber stabilized at 37°C under 5% CO2. Single channel live imaging was performed on an Axio oberver7 inverted Zeiss microscope using a 63× PlanApo oil objective (numerical aperture = 1.4) and captured using a Hamamatsu Orca-fusion 2 cameras through a mCherry filter set. We used a mercury lamp (Osram HBO 100 W/2). Images were taken distally to the axon initial segment (AIS) for 50s with an exposure time of 150 to 160 ms per image Zen blue 2.6 software (Zeiss).

#### Confocal microscopy

Confocal microscopy was performed using an upright FV-1200 Confocal Microscope and an inverted SP8 Leica confocal microscope, with 40x or 63x oil immersion objectives (1.30 and 1.40 numerical aperture respectively), controlled by Metamorph (FV-1200) or LasX (SP8, Leica, v 3.5.6) softwares. To test the effect of neuronal activity on node-like cluster formation for each acquisition, 387.69 μm x 387.69 μm of 2048×2048 pixels image stacks, with a z-step of 0.35 μm were acquired with the 40x objective using 405, 488, 565 and 647 laser lines. For imaging of the organotypic culture slices, fields of view with Purkinje cells expressing whether mCherry (for chemogenetic experiment) or YFP (for optogenetics experiment) were chosen. A stack sufficient to follow their axons was taken. When imaging to quantify the density of node like clusters a minimal field of view of 123.14 μm x 123.14 μm of 1024×1024 pixels image stack, including at least 10 Z-series with a z-step of 0.30 μm was acquired with a 63x objective.

In order to quantifiy the density of node like clusters in mouse spinal cord, we imaged fields in the center of the LPC lesion (5 images per animal). Each acquisition corresponds to a 160.61 μm x 160.61 μm image stack (1024×1024 pixels), including at least 20 images per stack, with a z-step of 0,30 μm. All the images were acquired with an inverted SP8 Leica microscope with a 405, 488, 565 and 647 laser line and a 63x oil immersion objective with a numerical aperture of 1.40, controlled by LasX software (SP8, Leica, v 3.5.6).

### Analysis

#### In vitro axonal transport analysis

For the analysis of axonal transport live imaging in vitro, kymographs (representing the trajectory of the fluorescent puncta over time) were generated with the ImageJ software, using the KymoToolBox plugin (kindly provided by Dr F. Cordelière, University of Bordeaux, France). For analysis, individual trajectories were manually traced. For all kymographs, moving puncta [anterograde (toward the axon terminal), retrograde (toward the cell body), or bidirectional] were defined as previously described^9^. We calculated the mean number of puncta per 100 μm in each condition and the distribution of anterograde, retrograde, bidirectional, and stationary puncta. Group analysis of the trajectories was performed using a homemade Excel macro designed by Dr J. Tailleur (University of Paris Diderot, France and MIT, Cambridge, USA). The data presented correspond to the analysis of 17 to 33 neurons, from 3 to 4 experiments. Data are presented as the mean ± standard error of the mean (SEM) of the experiments.

#### Study of nodal marker expression in GABAergic neurons in vitro

GABAergic neurons were colabelled for GAD67, PanNav and a nodal marker of interest (β1Nav, β 2Nav, Nfasc, Nav1.1, Kv3.1b or KCNQ2/Kv7.2) at 17 DIV, following KYN treatment or in control condition. The number of GAD+ neurons with node-like clusters (defined by PanNav expression) using an upright Axio imager microscope (Zeiss) with a 40x oil immersion objective with a numerical aperture of 1.3. A Nav cluster was defined by a length between 1 and 8 μm and a neuron was counted as positive for node like clusters when at least 2 Nav clusters were observed along its axon ^8^. The expression of the tested nodal marker of interest was assessed in NLC+ and NLC-GAD+ populations. A total of about 250 to 600 GAD+ neurons were counted per condition, from three to four independent experiments (biological replicates). Results are expressed as mean ± SEM of the experiments.

#### Quantification of node-like clusters

For hippocampal mixed cultures, miRNA transfected or DREADDS expressing GABAergic neurons were labelled for GAD67 and GFP or mCherry respectively. The number of GAD67+/GFP+ or mCherry+ neurons with node-like clusters was quantified as described above. On average about 60-200 GAD+ or GAD+/GFP+ or mCherry+ neurons were counted per experiment per condition, and at least three independent experiments were performed (biological replicates). Results are expressed as mean ± SEM.

For the cerebellar slice cultures at 3-4 DIV and for the cerebellum sections at P10, the density of node-like clusters was quantified on one optical section in the middle of the image stack. The rest of the stack was used to make sure that the structures corresponded to a node-like cluster and not to an axon initial segment orthogonal to the imaging plan. The quantification was performed on 5 different fields of view per animal, the number of node-like clusters per area unit was averaged to give the mean value for one animal. The analysis was performed in at least 5 animals per condition from at least two different independent litters.

Following neuronal activity stimulation with both DREADDs and optogenetics, the proportion of Purkinje cells with node-like clusters, the number of node-like clusters was assessed along the first 130 μm of the axon starting from the distal end of the axon initial segment. When the axon had a branch point along these 130 μm, the analysis was made along the main branch going toward the white matter tract. The analysis was reduced to this part of the axon to restrain the quantification to the area where myelination was ongoing. Indeed, during cerebellar development the myelination proceeds in an ascending manner along Purkinje axons from the white matter tract toward the soma ^74^. The proportion of Purkinje cells with node like clusters were quantified on 4 to 5 different fields of view to obtain on average 32 (range: 16-48) mCherry+ or YFP+ Purkinje cells per condition per animal for myelinating condition and on average 31 (range: 21-38) for remyelination.

To analyze the effect of neuronal activity on node like clustering in vivo during remyelination, the number of node-like clusters, identified as isolated Nav clusters (i.e. without Caspr staining surrounding the cluster) were quantified on a central image of the image stack, the surrounding images further being used to exclude the possibility of the structures being heminodes or mature nodes. For each image, the density of node-like clusters was calculated as the number of clusters per image area, brought back to number of clusters per mm^2^ and the mean density was calculated using the 5 images acquired per animal. The results are presented as the mean number of node-like clusters per area per animal (n=7 animals hM3/NaCl, n=6 animals hM3/CNO, n=6 animals hM4/NaCl and n=5 animals hM4/CNO).

#### Statistics

All statistical analysis and data visualization were carried out using Prism (GraphPad, version 7) and R software version 3.6.2. For all the experiments, the number of biological replicates and statistical tests applied are reported in the text or in the figure legends, as well as in the summary Table. Graphs and data in the text are reported as the mean ± SEM, each biological replicate is individually plotted. The level of statistical significance was set at p < 0.05 for all tests.

When the sample size was n ≥ 6, we assessed for the normality of the distribution using a Shapiro– Wilk test and used accordingly parametric tests when the distribution was not significantly different than a normal distribution and non-parametric tests were applied otherwise. When the sample size was n<6, parametric tests were used assuming the normality of the distributions since in these cases each individual sample represented a large number of repeated measures.

For the in vitro studies, Student’s t-test or One– or Two-way ANOVA followed by multiple comparison tests were used when appropriate. For the analysis of the effect of neuronal activity on node-like clustering *ex vivo,* the design involved a pairing, therefore groups were compared using two-sided paired tests. For the quantification of node-like clusters density *in vivo* and *ex vivo*, two-sided unpaired tests were used.

All the statistical results are summarized in the supplementary Table 1.

## Acknowledgments

AD, NSF, RR, CP designed the research; RR, CP, EM, MT, MS, PS, NSF and AD did the experiments and analysis; FX Lejeune participated in statistical analysis; AD, NSF, RR, CP, BS and CL wrote the publication. We thank Eric Marty (Phymep, INSERM) and Julien Ballbe y Sabate for their help in generating the optogenetics set-up; Boris Zalc and Julien Tailleur for their critical reading of the paper; Clement Lena for kindly providing the L7Chr2-YFP mouse line. We thank the ICM platforms implicated in this project: iQuant and David Akbar, EPhys and Charlotte Deleuze, PhenoMice, Celis, the histology platform and iGenSeq. This work was funded by ARSEP grants to AD and NSF, BBT-IHU grant to AD and ICM and INSERM support. RR and MT present addresses are Department of Neuromuscular Diseases, University College London, London, United Kingdom and Zebrafish Neurogenetics Unit, Institut Pasteur, CNRS UMR3738, Université de Paris Cité, Paris, France.

