## Supplementary_data for "Neuronal activity promotes axonal node-like clustering prior to myelination and remyelination in the central nervous system"

SUPPLEMENTARY FIGURE LEGENDS

**Figure S1. The inhibition of glutamatergic receptors affects some nodal protein expression, but does not impact nodal marker axonal transport.**

(A-C) Expression (+ or -) of Nav1.1 (A), β1Nav (B) and Kv3.1b (C), in GABAergic neurons with or without clusters (NLC + or – respectively) in control condition (black bars) or following kynurenic acid treatment (red bars). (D) Representative kymographs illustrating β1Nav- and β2Nav-mCherry axonal transport at 17 DIV following kynurenic acid treatment (KYN) compared to control condition. Anterograde transport from left to right. (E-H) Quantification of the total number of mCherry+ puncta per 100 μm (E, G) and mCherry+ puncta category distribution (F, H) for β1Nav- and β2Nav-mCherry following KYN (red bars) treatment compared to control (black bars). FO: forward, BA: backward and BI: bidirectional moving puncta. ST: stationary puncta. The histograms show the means ± SEM. (E-G) Unpaired t test, (F, H) Two-way ANOVA followed by Tukey's multiple comparisons test. β1Nav-mCherry: n=4 experiments; β2Nav-mCherry: n=3 experiments. (A) Scale bar: 5s.

**Figure S2. Node-like clusters are formed prior to myelination and remyelination along Purkinje cells axons.** (A) Immunohistostainings on a cerebellar organotypic slices at 4 DIV showing node-like clusters (Nav in red, filled arrowheads) without paranodal clustering (gray, Caspr) distributed in regions with ongoing myelination (PLP, in green). The diffused Caspr signal allow to follow unmyelinated Purkinje axons. (B) Example of a node-like cluster (Nav in red, filled arrowhead) isolated surrounded by two heminodes (contour arrowheads) along the same axons (dashed line traced using the faint diffuse Caspr staining along the axon). (C) Orthogonal projections showing isolated node-like clusters (Nav in red, filled arrowhead) without paranodal clustering (gray, Caspr) in a remyelinating (PLP, in green) cerebellar slice at 11 DIV. (D) Orthogonal projection of a sagittal section of the cerebellum at P10 showing Purkinje cells axons (Calbindin, in gray), with node-like clusters (Nav in red, white filled arrowhead) along unmyelinated part of their axons (myelin stained by PLP, green). (E) Orthogonal projection showing at a higher magnification two node-like clusters (Nav in red, filled arrowhead) along an unmyelinated portion of the same axon and heminodes (contour arrowhead) at the extremity of the myelin sheathes. (F) Quantifications of node-like clusters in the cerebellum *in vivo* at P10, *ex vivo* at 3-4 DIV and *ex vivo* in remyelinating area at 11 DIV show similar densities of node-like structures in the regions with ongoing myelin deposition. Each value individually plotted corresponds to 1 animal, *in vivo* P10 and *ex vivo* 11 DIV: n=4 animals, *ex vivo* 4 DIV: n=5 animals. (F) One-way ANOVA. Scale bars: (A, C) 20 μm, (B,D) 5 μm, (D) 10 μm.

**Figure S3. Modulations of the firing activity of Purkinje cells in organotypic cerebellar slices by DREADD and optogenetic stimulations.** (A) Example of a hM3D(Gq)-mCherry transduced Purkinje cell (filled white arrowhead) recorded in loose cell-attached voltage clamp. (B) Representative example of loose-cell attached voltage clamp recordings on a hM3D(Gq)-mCherry transduced Purkinje cells, in control condition (first raw) followed by CNO treatment (0.5 μM, second raw). (C) Quantification of the mean firing frequency of hM3D(Gq)-mCherry transduced Purkinje cells in control condition (CTRL) and following addition of CNO. (D, E) Immunohistochemistry showing the expression of ChR2-YFP (in green) specifically in Purkinje cells (Calb positive, in red) at 4 DIV (D) and 10 DIV (E). (F) Quantification of the percentage of Purkinje cells expressing YFP in folia of the cerebellum with high density of YFP signal (folia usually used for all the analysis performed on organotypic slices). (G) Example of a ChR2-YFP expressing Purkinje cell (in green, filled white arrowhead) recorded in loose cell-attached voltage clamp. (H) Representative examples of loose-cell attached voltage clamp recordings on a ChR2-YFP expressing Purkinje cells in control slices, without optogenetic stimulation (LED off, first raw) followed by stimulation at 10Hz with 10ms long pulses at 1,5mW/mm$^2$ (second raw). The pattern of neuronal firing (in black) follows the pattern of light (pulses are indicated with the blue rectangles). (I) Quantification of the mean firing frequency of Purkinje cells without optogenetic stimulation (CTRL) and following optogenetic stimulation (ACT) in myelinated slices. (C, I) Wilcoxon matched-pairs signed rank test. Each individual point represents the mean for one cell recorded. n=8 cells from 4 animals (C) and n=6 cells from 4 animals (I). (F) n=3 animals per condition. Scale bars: (D, E) 30 μm.

**Figure S4. The inhibition of glutamatergic transmission decreases node-like cluster formation ex vivo during myelination.**

(A) Immunostaining of cerebellar slices showing Purkinje cells (Calbindin, blue) with node like-clusters (Nav, red, not associated to myelin, PLP, white). Following kynurenic acid treatment (KYN, 1mM), fewer Purkinje cells assemble node-like clusters compared to control condition. Node-like clusters are shown with white filled arrowhead and the corresponding Purkinje cell expressing node-like cluster are shown with white contour arrowhead. (B) Quantification of the percentage of Purkinje cells with node-like clusters following kynurenic acid treatment (KYN). The slices were fixed at 3 DIV at the onset of myelination. Each point corresponds to one animal. n=6 animals per condition. Paired t test. (B) Scale bar: 20 μm.

**Figure S5. Validation of the in vivo DREADDs approach coupled to focal demyelination of mouse spinal cord**

(A) Illustration of mouse motor cortex transduced with AAVrg-hSyn-hM3Gq-mCherry showing cells efficiently expressing the hM3Gq receptor (mCherry, red). (B) Mouse cortical neurons transduced with

AAVrg-hSyn-hM3D(Gq)-mCherry (left panel) or AAVrg-hSyn-hM4D(Gi)-mCherry (right panel) showing an increase or a decrease of cFos expression (in green) in neurons expressing hM3D(Gq) or hM4D(Gi) respectively (mCherry, red) following CNO injection compared to control. mCherry+ cells expressing cFos are shown with white filled arrowhead. (C-D) Quantification of the percentage of mCherry+ neurons expressing cFos in mouse transduced with AAVrg-hSyn-hM3D(Gq)-mCherry (C) or AAVrg-hSyn-hM4D(Gi)-mCherry (D). Each point corresponds to one animal. (C) n = 4 animals, Mann-Whitney test. (D) n = 4 animals for NaCl condition and n=6 for CNO condition, Mann-Whitney test. (A-B) Scale bar: (A) 1mm; (B) left panel: 30 µm, right panel: 20µm.

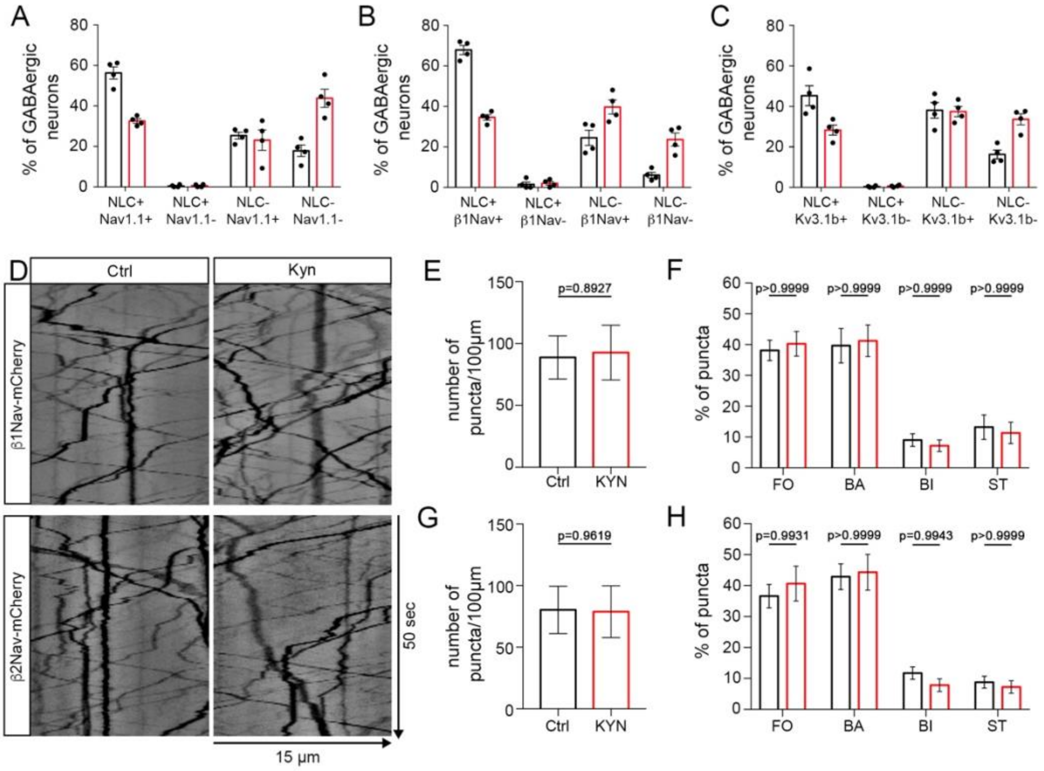

A
B
C
% of GABAergic neurons
NLC+ Nav1.1+
NLC+ Nav1.1-
NLC- Nav1.1+
NLC- Nav1.1-
NLC+ β1Nav+
NLC+ β1Nav-
NLC- β1Nav+
NLC- β1Nav-
NLC+ Kv3.1b+
NLC+ Kv3.1b-
NLC- Kv3.1b+
NLC- Kv3.1b-
D
Ctrl
Kyn
β1Nav-mCherry
β2Nav-mCherry
50 sec
15 µm
E
G
number of puncta/100µm
p=0.8927
p=0.9619
Ctrl
KYN
F
H
% of puncta
p>0.9999
p=0.9931
p=0.9943
FO
BA
BI
ST


Figure S1

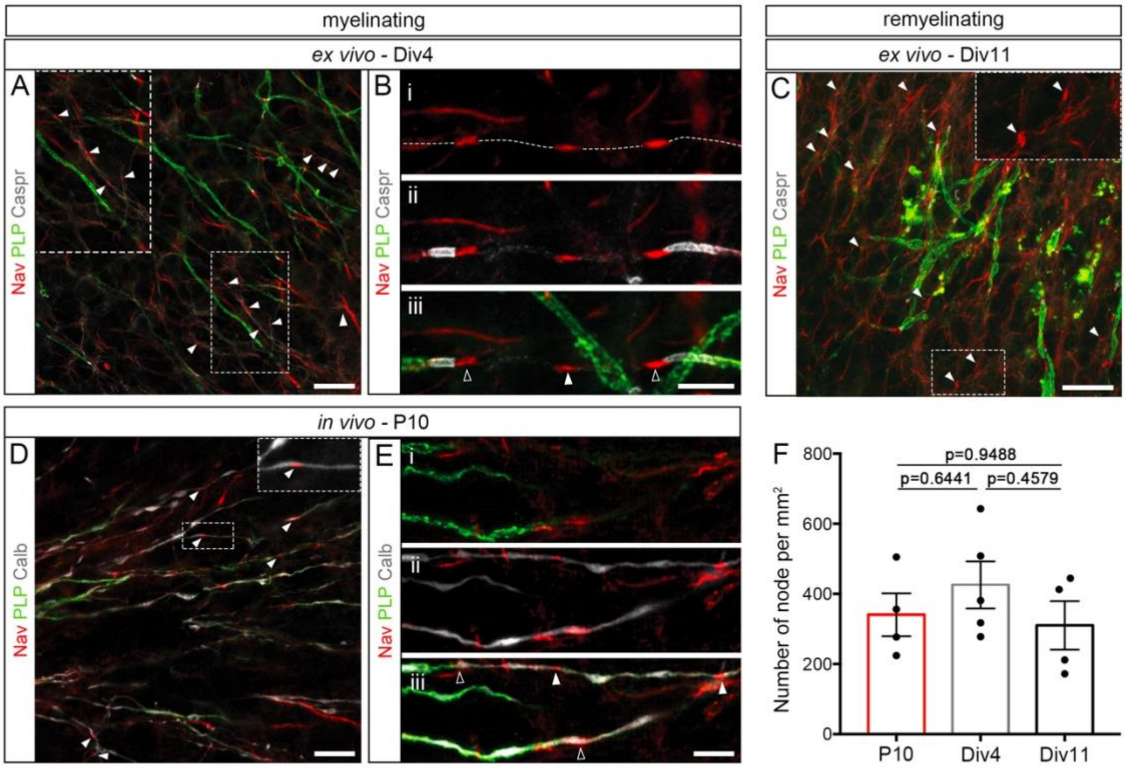

myelinating
remyelinating
ex vivo - Div4
ex vivo - Div11
A
B
C
Nav PLP Caspr
i
ii
iii
in vivo - P10
D
E
Nav PLP Calb
F
Number of node per mm²
800
600
400
200
0
p=0.9488
p=0.6441
p=0.4579
P10
Div4
Div11


Figure S2

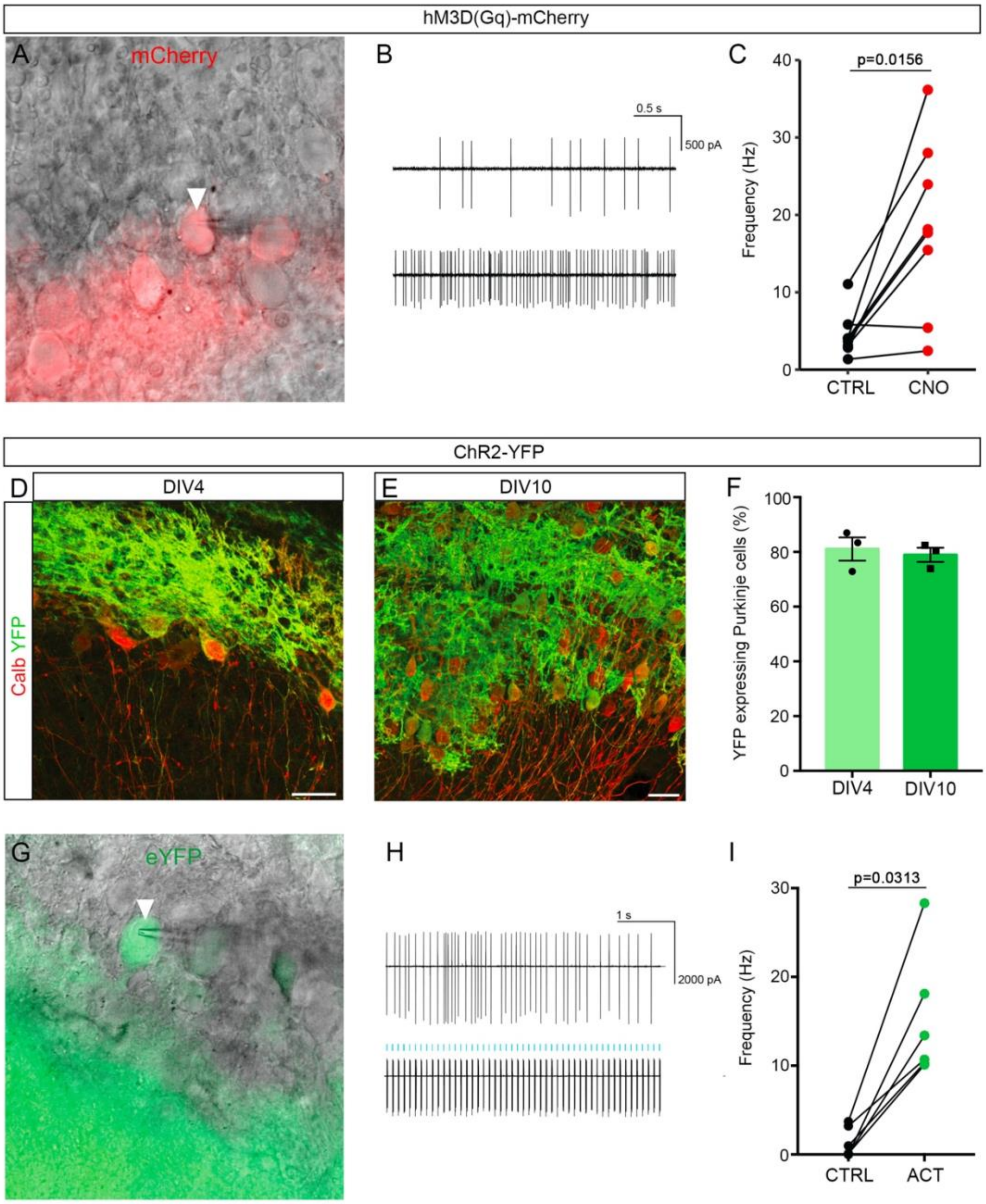
hM3D(Gq)-mCherry
A
mCherry
B
0.5 s
500 pA
C
p=0.0156
Frequency (Hz)
CTRL
CNO
ChR2-YFP
D
DIV4
Calb YFP
E
DIV10
F
YFP expressing Purkinje cells (%)
DIV4
DIV10
G
eYFP
H
1 s
2000 pA
I
p=0.0313
Frequency (Hz)
CTRL
ACT

Figure S3

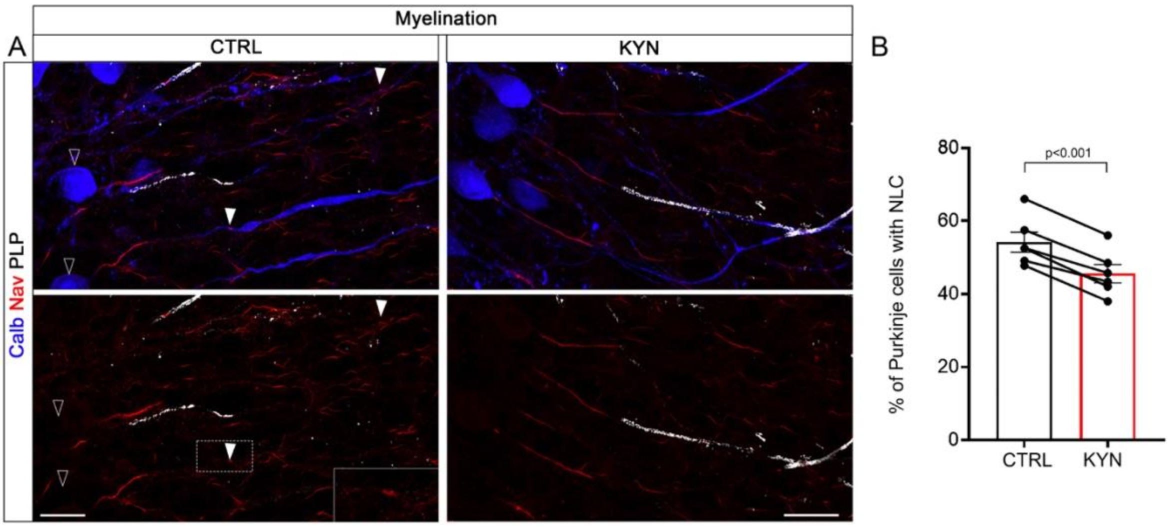
Myelination
A
CTRL
KYN
B
Calb Nav PLP
% of Purkinje cells with NLC
p<0.001
80
60
40
20
0
CTRL
KYN

Figure S4

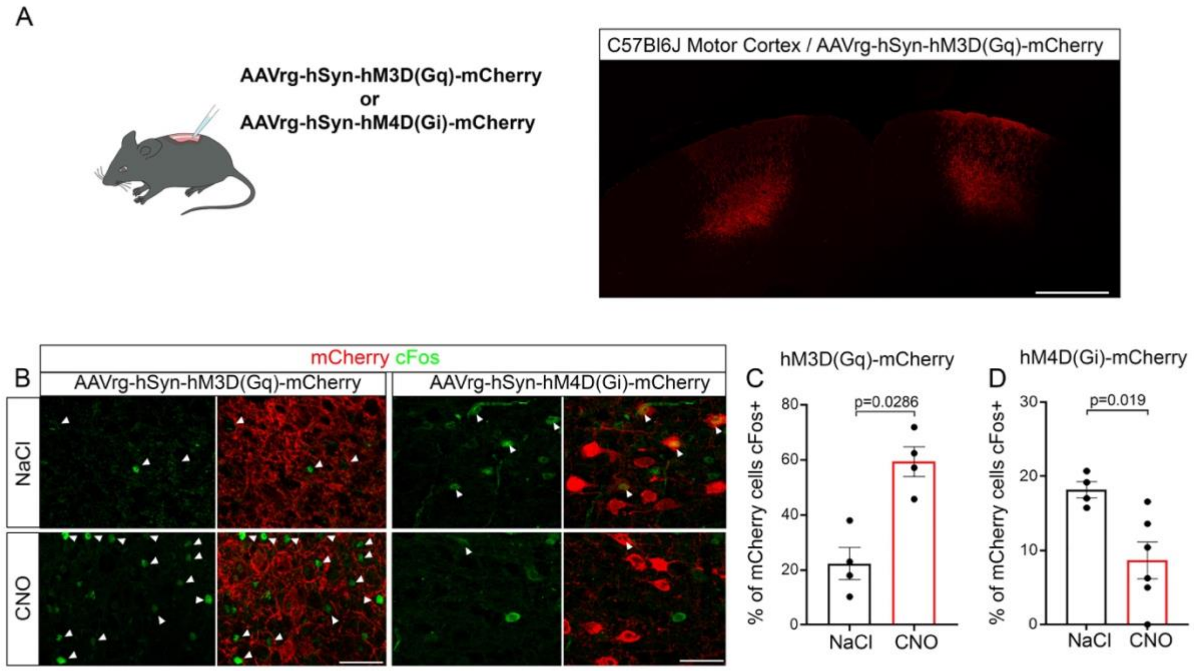

A
AAVrg-hSyn-hM3D(Gq)-mCherry
or
AAVrg-hSyn-hM4D(Gi)-mCherry
C57Bl6J Motor Cortex / AAVrg-hSyn-hM3D(Gq)-mCherry
B
mCherry cFos
AAVrg-hSyn-hM3D(Gq)-mCherry
AAVrg-hSyn-hM4D(Gi)-mCherry
NaCl
CNO
C
hM3D(Gq)-mCherry
p=0.0286
% of mCherry cells cFos+
0
20
40
60
80
NaCl
CNO
D
hM4D(Gi)-mCherry
p=0.019
% of mCherry cells cFos+
0
10
20
30
NaCl
CNO


Figure S5

**SUPPLEMENTARY TABLE 1**

| Experience | Figure number | Exact value | Sample | Test used | P value |
|---|---|---|---|---|---|
| **Neuronal activity impact on the formation of node-like clusters in vitro hM4D(Gi)** | Figure 1C | **mean ± SEM: ctrl = 46.90 ± 4.93% ; CNO = 40 .10 ± 4.74%** | n=3 independent experiments (treated vs non treated GABAergic mCherry positive neurons) | Paired t test | % of GABAergic neurons with node-like clusters: ctrl vs. CNO: p = 0.0011 |
| **Neuronal activity impact on the formation of node-like clusters in vitro hM3D(Gq)** | Figure 1D | **mean ± SEM: ctrl = 46.10 ± 6.27% ; CNO = 58.00 ± 7.61%** | n=4 independent experiments (treated vs non treated GABAergic mCherry positive neurons) | Paired t test | % of GABAergic neurons with node-like clusters: ctrl vs. CNO: p = 0.0483 |
| **Node like clustering in mixed cultures treated with TTX/KYN /NBQX /APV** | Figure 1F | **mean ± SEM : Ctrl = 37.03 ± 2.09; TTX = 11.60 ± 1.86 ; KYN = 19.40 ± 1.99; NBQX = 22.87 ± 2.40 ; APV = 31.93 ± 1.57** | n = 4 independent experiments for Ctrl and TTX and n = 3 for KYN/NBQX/APV | One-Way ANOVA followed by Dunnett's multiple comparison test | % of GAD Neurons with NLC : Ctrl vs TTX: p < 0.0001 ; Ctrl vs KYN: p = 0,0002 ; Ctrl vs NBQX: p = 0.0012 ; Ctrl vs APV: p= 0.2812 |
| **Node like clustering on purified neurons + OCM treated with TTX/KYN/NBQX/APV** | Figure 1H | **mean ± SEM : Ctrl = 33.67 ± 1.07 ; TTX = 10.93 ± 0.79; KYN = 13.77 ± 0.39; NBQX = 19.30 ± 1.78; APV = 14.53 ± 1.28** | n = 3 independent experiments | One-Way ANOVA followed by Dunnett's multiple comparison test | % of GAD Neurons with NLC : Ctrl vs TTX: p < 0.0001; Ctrl vs KYN: p < 0.0001; Ctrl vs NBQX: p < 0.0001; Ctrl vs APV: p < 0.0001 |
| **Nodal marker expression in mixed cultures treated with KYN** | Figure 2 | **mean ± SEM : β1Nav Ctrl = 92.39 ± 1.70; KYN = 67.70 ± 3.10; Nav1.1 Ctrl = 81.70 ± 2.86; Kyn = 55.65 ± 4.45; Kv3.1b Ctrl = 83.38 ± 1.95; KYN = 65.75 ± 2.54; Kv7.2 Ctrl = 99.37 ± 0.63; KYN = 99.00 ± 1.00; Nfasc Ctrl = 98.80 ± 1.20; KYN = 98.77 ± 1.23; β2NAv Ctrl = 95.85 ± 1.69; KYN = 95.38 ± 1.63** | b1Nav: Ctrl :351 cells; KYN: 325 cells, n=4 independent experiments. Nav1.1: Ctrl : 558 cells; Kyn : 530 cells, n=4 independent experiments. Kv3.1b: Ctrl: 582 c ells; KYN: 622, n=4 independent experiments. Kv7.2: Ctrl : 366 cells; Kyn : 371 cells, n=3 independent experiments. Nfasc: Ctrl: 246 cells, KYN: 251 cells, n=3 independent experiments. b2Nav: Ctrl: 333 cells; KYN: 434 cells, n=3 independent experiments. | Unpaired t-test | % of GAD expressing thenodal marker of interest : Ctrl vs KYN: β1Nav: p = 0.0022 ; Nav1.1: p = 0.0026 ; Kv3.1b: p = 0.0015 ; Kv7.2: p = 0.7722; Nfasc: p = 0.9855; β2NAv: p = 0.8463 |
| **Prenodal clustering in miRCtrl vs miRNav1.1 254 vs miRNav1.1 947** | Figure 3 | **mean ± SEM : panNav miRCtrl = 60.07 ± 1.82; miR 254 = 32.88 ± 1.36; miR 947 = 23.38 ± 2.94; AnkG miRCtrl = 62.66 ± 1.76; miR 254 = 24.67 ± 1.29; miR 947 = 21.40 ± 1.21** | PanNav : n=3-4 independent experiments AnkG: n=4-5 independent experiments | One-Way ANOVA followed by Dunnett's multiple comparison test | % of GAD Neurons with NLC : AnkG: miRCtrl vs miR254: p < 0.0001 ; miRCtrl vs miR947: p < 0.0001 ; **panNav:** miRCtrl vs miR254: p < 0.0001 ; miRCtrl vs miR947: p < 0.0001 |
| **Neuronal activity impact on node-like clustering along Purkinje cells ex vivo prior to myelination (hM3D(Gq))** | Figure 4D | **mean ± SEM: Ctrl = 39.66 ± 2.89% ; CNO = 56.17 ± 3.28%** | n=5 animals (DMSO vs CNO treated slices from the same animal) | Paired t test | % of mCherry+ Purkinje cells with node-like clusters: p = 0.0171 |
| **Neuronal activity impact on node-like clustering along Purkinje cells ex vivo prior to myelination (Optogenetics)** | Figure 4G | **mean ± SEM: Ctrl = 43.03 ± 1.79% ; opto=59.97 ± 4.39%** | n=5 animals (ctrl vs opto slices from the same animal) | Paired t test | % of YFP+ Purkinje cells with node-like clusters: p = 0.0058 |
| **Neuronal activity impact on node-like clustering along Purkinje cells ex vivo prior to remyelination (hM3D(Gq))** | Figure 5D | **mean ± SEM: Ctrl = 40.60 ± 2.52% ; CNO = 51.60 ± 2.57%** | n=6 animals (DMSO vs CNO treated slices from the same animal) | Paired t test | % of mCherry+ Purkinje cells with node-like clusters: p = 0.0225 |
| **Neuronal activity impact on node-like clustering along Purkinje cells ex vivo prior to remyelination (Optogenetics)** | Figure 5G | **mean ± SEM: Ctrl = 49.07 ± 1.89% ; opto =58.89 ± 2.72%** | n=6 animals (ctrl vs opto slices from the same animal) | Paired t test | % of YFP+ Purkinje cells with node-like clusters: p = 0.0038 |
| **Neuronal activity impact on node like-cluster density in vivo prior to remyelination (hM3D(Gq))** | Figure 6C | **mean ± SEM : Ctrl = 652 ± 129; CNO = 907 ± 44** | n=7 animals for Ctrl (NaCl) and n=6 animals for CNO | Mann-Whitney test | Number of node like-clusters per mm2: p = 0.0350 |
| **Neuronal activity impact on node like-cluster density in vivo prior to remyelination (hM4D(Gi))** | Figure 6E | **mean ± SEM : Ctrl = 914 ± 69 ; CNO = 630 ± 44** | n=7 animals for Ctrl (NaCl) and n=5 animals for CNO | Mann-Whitney test | Number of node like-clusters per mm2: p = 0.0087 |
| **SUPPLEMENTARY FIGURES** | | | | | |
| **Modulation of nodal marker axonal transport by KYN : Number of puncta/100 µm** | Figure S1E, G | **mean ± SEM: β1NavCtrl = 89 ± 17 ; KYN = 93 ± 22; β2Nav Ctrl = 80 ± 19 ; KYN = 79 ± 21** | β1Nav : n=4 independent experiments β2Nav: n=3 independent experiments | Unpaired t test | Number of puncta/100µm: β1Nav: p=0.8927; β2Nav: p=0.7000 |
| **Modulation of nodal marker axonal transport by KYN : puncta distribution (%; FO: forward, BA: backward, BI: bidirectionnal: ST: stationary)** | Figure S1F, H | **mean ± SEM: β1Nav Ctrl: FO = 38.1 ± 3.3; BA = 39.7 ± 5.6; BI = 9.0 ± 2.0; ST = 13.2 ± 4.0; β1Nav KYN: FO = 40.3 ± 4.0; BA = 41.2 ± 5.16; BI = 7.2 ± 1.9; ST = 11.3 ± 3.5; β2Nav Ctrl: FO = 36.6 ± 3.8; BA = 42.9 ± 4.1; BI = 11.7 ± 2.0; ST = 8.8 ± 1.9; β2Nav KYN: FO = 40.6 ± 5.6; BA = 44.3 ± 5.8; BI = 7.8 ± 2.1; ST = 7.2 ± 2.0** | β1Nav : Ctrl = 24 movies; KYN = 33 movies, from n=4 independent experiments β2Nav: Ctrl = 17 movies; KYN = 23 movies, from n=3 independent experiments | Two-way ANOVA followed by Tukey's multiple comparisons test | % of puncta per category: Ctrl vs KYN: β1Nav: all categories p > 0.999; β2Nav: FO p = 0.9931; BA p > 0.9999; BI p = 0.9943; ST p > 0.9999 |
| **Node-like cluster density in vivo (myelination, cerebellum) vs ex vivo (myelination) vs ex vivo (remyelination)** | Figure S2F | **mean ± SEM: in vivo = 340 ± 61 per mm2 ; ex vivo = 426 ± 67 per mm2; remyelination = 310 ± 69 per mm2** | n=4 animals (in vivo); n= 5 animals (ex vivo myelination); n=4 animals (ex vivo remyelination) | One-way ANOVA followed by Tukey's multiple comparisons test | Number of node-like clusters per mm2: in vivo vs. ex vivo myelination: p = 0.6441; ex vivo myelination vs. ex vivo remyelination: p = 0.4579 ; in vivo myelination vs. ex vivo remyelination: p = 0.9488 |
| **Firing frequency of hM3D(Gq) positive Purkinje cells in Ctrl vs CNO treated condition ex vivo** | Figure S3C | **mean ± SEM: ctrl = 4.5 ± 1.0 Hz ; CNO = 18.4 ± 4.0 Hz** | n=8 cells from 4 animals | Wilcoxon paired test | Firing frequency: p = 0.0156 |
| **Percentage of YFP+ Purkinje cells ex vivo** | Figure S3F | **mean ± SEM: Div4 = 81.06 ± 4.23% ; Div10 = 78.93 ± 2.58%** | n=3 animals (Div4); n=3 animals (Div10) | no test | no test |
| **Firing frequency of YFP+ Purkinje cells in Ctrl vs 10Hz stimulation condition ex vivo** | Figure S3I | **mean ± SEM: ctrl = 1.3 ± 0.7 Hz ; opto = 15.2 ± 2.9 Hz** | n=6 cells from 4 animals | Wilcoxon paired test | Firing frequency: p = 0.0313 |
| **Impact of glutamatergic input inhibition on node like-clustering ex vivo prior myelination** | Figure S4B | **mean ± SEM : Ctrl = 54,24 ± 2,727 ; KYN = 45,60 ± 2,533** | n=6 animals (Ctrl vs KYN treated slices from same animal) | Paired t test | % of Purkinje cell with node like cluster: p < 0,0001 |
| **Validation of neuronal activity modulation (cFos expression) in vivo (mice transduced with hM3D(Gq))** | Figure S5C | **mean ± SEM : NaCl = 22.32 ± 5.85; CNO = 59.29 ± 5.45** | n= 4 animals per condition | Mann-Whitney test | % of mCherry+ cells expressing cFos: p = 0,0286 |
| **Validation of neuronal activity modulation (cFos expression) in vivo (mice transduced with hM4D(Gi))** | Figure S5D | **mean ± SEM : NaCl = 18.19 ± 1.11; CNO = 8.65 ± 2.48** | n=4 animals for Ctrl and 6 for CNO condition | Mann-Whitney Test | % of mCherry+ cells expressing cFos: p = 0,019 |
